# Unlocking substrate specificities of human solute carrier proteins using untargeted metabolomics

**DOI:** 10.64898/2026.07.27.740914

**Authors:** Yuesheng Zhang, Lyubomir Dimitrov Stanchev, Felicia Cara Schulz, Daniela Rago, Carlos G. Acevedo-Rocha, Alberto Santos Delgado, Douglas B. Kell, Irina Borodina

## Abstract

The limited understanding of transporter substrate spectra constrains our ability to interpret cell and membrane function, highlighting the need for methods that enable transporter deorphanization and characterization of promiscuous transport activities. Here, we present a *Xenopus* oocyte-based platform for unbiased transporter substrate discovery. Oocytes expressing heterologous solute carrier proteins (SLC) were incubated in human blood serum, a chemically complex metabolite library containing thousands of endogenous metabolites and xenobiotics, followed by paired untargeted LC-MS/MS profiling of intracellular extracts and surrounding medium to capture metabolite exchange events. Across the five human SLC transporters, viz. SLC10A2, SLC10A6, SLC13A2, SLC16A10, and SLC46A1, metabolite exchange signatures were detected, and automated feature annotation was refined by manual chromatographic peak inspection. The workflow recovered known substrates of SLC10A2 and SLC16A10 and identified additional transported metabolites with MS/MS confirmation. This method provides a scalable framework for transporter substrate profiling and prioritization of candidates for targeted validation.

## 1. Introduction

Transport of small molecules across cellular membranes is fundamental to cell physiology, governing the uptake and efflux of nutrients, metabolic intermediates, signaling molecules, and xenobiotics. These essential functions make membrane transporters attractive targets both for therapeutic development and for engineering improved cell factories [1]. Inhibitors of solute carriers (SLCs) have been used to develop cancer therapies and modulate the cellular uptake or disposition of metabolites and xenobiotics [1–3]. Transporter engineering has enabled cell factories to increase the productivity of converting cheap carbon substrates into secreted high-value products [5]. However, advancement in the field is hindered by the incomplete functional annotation of many transporters, highlighting the need for deorphanization [6]. Most small molecules interact with multiple proteins [7], with transporter promiscuity remaining a major challenge. Many SLCs, in addition to their roles in the transport of endogenous metabolites, are also involved in the transport of xenobiotics such as pharmaceuticals and antioxidants [8]. Similarly, in microbial cells, plant natural products can traffic in and out of engineered microbial cell factories even though no uptake transporter has been identified [9]. These observations suggest that many transporters exhibit broader and context-dependent substrate ranges than expected, while non-intuitive substrates are rarely explored in conventional characterization efforts.

Such knowledge gaps underscore the need for technologies capable of resolving transporter substrate spectrum. Traditional phenotypic assays, while robust for identifying transporters linked to growth-coupled traits, are difficult to extend to non-growth-coupled substrates that do not yield discernible phenotypes [10]. Classical transport assays, ranging from radiolabeled uptake and flux measurements to electrophysiological recordings, typically interrogate one substrate at a time. Although these methods provide high sensitivity and detailed kinetic information, they are inherently low-throughput and offer limited ability to uncover unexpected substrates or to capture substrate competition and inhibition effects.

To overcome these limitations, one strategy involves simply exposing cells expressing transporters of interest to media (such as human serum) containing potentially thousands of candidate metabolite substrates and assessing differential uptake by methods such as mass spectrometry [11–14]. This can be done by analysis of both extracellular and intracellular metabolomes, the former being considerably more convenient.

More recently, Møller-Hansen et al. expressed yeast transporters in *Xenopus* oocytes to generate a SLC gene library and screened their activity against six biotechnologically relevant metabolites, revealing notable differences in substrate promiscuity among the transporters [10]. Within this framework, an important next step is to expand the diversity and size of substrate libraries to enable higher-throughput, unbiased detection of transport activities. Human blood serum provides an attractive option: its chemically rich composition, when combined with untargeted metabolomics, allows thousands of metabolites to be surveyed in a single measurement. As mentioned above, Radi *et al*. demonstrated such an approach by using serum-based untargeted metabolomics to deorphanize transporters in *Escherichia coli* [13]. Harnessing 26 paired knockout and overexpression *E. coli* transporter mutants, the authors used mutant-dependent metabolic fingerprints to identify previously unrecognized transporter-substrate interactions.

Although microbial hosts such as bacteria and yeast are commonly used to study transporters, their strong native metabolism and many endogenous transporters can interfere with transport assays and make it harder to isolate the effect of the transporter being tested. *Xenopus* oocytes, by contrast, contain a minimal repertoire of native transporters, offering a cleaner background. They have been widely adopted as a versatile platform for characterizing transporters from both prokaryotic and eukaryotic origins [15,16]. In previous work, we demonstrated this advantage by profiling yeast amino acid transporters using targeted LC-MS/MS with ¹³C-labeled yeast extract in the oocyte system, achieving higher accuracy and reproducibility compared with analogous assays performed in yeast cells [17].

In this study, we present a *Xenopus* oocyte-based workflow for mapping substrate ranges of membrane transporters. Five human solute carrier proteins were selected to represent diverse validated substrate classes and transport mechanisms, expressed in oocytes, and incubated with human blood serum. Metabolite exchange was quantified by untargeted metabolomics analysis of both extracellular serum and intracellular extracts. This approach revealed transporter-specific patterns of uptake and excretion. Importantly, these included metabolites consistent with previously reported activities, providing evidence for the accuracy of the method. In addition, we discovered several novel candidate substrates. The identified interactions were further evaluated *in silico* via molecular docking. This platform provides a generalizable strategy for elucidating the substrate specificities of potentially promiscuous membrane transporters across diverse biological systems.

## 2. Materials and methods

### 2.1. Materials

Cloning materials were obtained from New England Biolabs (Ipswich, MA, USA) or Thermo Scientific (Waltham, MA, USA). Gene sequencing service was provided by Eurofins Genomics (Ebersberg, Germany) and oligonucleotides were purchased from Integrated DNA Technologies (Leuven, Belgium). Synthetic genes for the human SLC transporters were ordered from Twist Bioscience with codon optimization for *Xenopus laevis* expression (CA, USA). All other materials were purchased from Sigma Aldrich (München, Germany), unless stated otherwise.

### 2.2. Cloning and plasmid construction

*E. coli* DH5α strain was used for plasmid amplification, and successful transformants were selected on LB plates with 100 mg/mL ampicillin. Cloning of transporter genes for cRNA expression in *Xenopus laevis* oocytes (Ecocyte Bioscience, Dortmund, Germany) was completed as previously described [23]. Briefly, transporters of the synthetic genes were first amplified by PCR and then cloned into the pCfB5245 plasmid encoding T7p-β-globin 5-UTR and β-globin 3-UTR. The resulting plasmids were then used as a PCR template for the amplification of the cassette fragment for *in vitro* RNA synthesis. Capped RNAs for transporter expression in oocyte cells were then synthesized with T7 mMESSAGE mMACHINE™ kit (Ambion). The quality of the cRNAs (> 500 ng/µL) was analysed by Agilent 2100 Bioanalyzer (Agilent Technologies) before proceeding with oocyte injection. All oligonucleotides and plasmids are listed in Supplementary Table S1 and S2, respectively.

### 2.3. Transport assay in oocyte cells

Transport assays in *Xenopus laevis* oocyte cells were performed as previously described [24] with the following modifications. Briefly, oocytes were injected with 50 nL of cRNA (400 ng/µL transporter cRNA and 100 ng/µL GFP cRNA) using an automated RoboInject device (Multichannel System, Germany) and incubated for three days at 18 °C. After 3 days of incubation, GFP positive cells were selected and used further for the transport assay.

Six to eight oocyte cells per biological replicate were pre-incubated in a Kulori buffer (90 mM NaCl, 1 mM KCl, 1 mM MgCl_2_, 1 mM CaCl_2_, 5 mM HEPES, pH 7.4) and then transferred to 50 µL human blood serum for 1 h at room temperature. For intracellular analysis, 40 µL of the blood serum were then transferred to a new Eppendorf tube and proteins precipitated by the addition of 140 µL acetonitrile. After centrifugation, 120 µL of the supernatant were transferred to a new tube and dried down for 3 h at 30 °C. The dried samples were then resuspended in 100 µL LC-MS grade water before LC-MS analysis. For extracellular analyses, the oocyte cells were transferred to 4 °C Kulori buffer to stop the assay and washed three times in fresh Kulori buffer before being disrupted with 100 µL ice-cold 50 % (v/v) methanol. Cell lysates were incubated for 2 h at –20 °C and then spun down to remove cell debris. Finally, 70 µL from the supernatant were mixed with 50 µL water before subjecting to LC-MS analysis.

### 2.4. Liquid chromatography mass spectrometry

The LC-MS/MS analyses were performed using an ultra-high-performance liquid chromatography (UHPLC) system (Vanquish Duo, Thermo Fisher Scientific, USA) coupled to an Orbitrap ID-X mass spectrometer (Thermo Fisher Scientific, USA) equipped with a heated electrospray ionization (HESI) source.

The chromatographic separation was achieved using a Waters ACQUITY BEH C18 column (2.1 mm × 100 mm, 1.7 μm) (WatersTM, Milford, MA, USA) maintained at 40 °C. The mobile phases consisted of LC-MS water (Honeywell, USA) containing 0.1% (v/v) LC-MS grade formic acid (VWR Chemicals, USA) (mobile phase A), and LC-MS grade acetonitrile (Honeywell, USA) containing 0.1 % (v/v) formic acid (mobile phase B) delivered at a flow rate of 0.35 mL min-1. The gradient elution program started at 2 % B, held for 0.8 min, followed by a linear increase to 5 % B over 3.1 min and to 100 % B over 6.7 min and held for 1 min. The column was then equilibrated for 2.7 min prior to the next injection. The injection volume was 1 µL.

The MS/MS measurements were acquired in positive-ion (ESI+) and negative-ion (ESI-) mode with a spray voltage of +3.5 kV and –2.5 kV, respectively. Full scan MS and data-dependent MS/MS spectra were acquired over an m/z range of 70-1,000 and in profile mode.

The MS¹ and MS² resolutions were set to 120,000 and 30,000, respectively. Data-dependent acquisition was performed using a method in which the most intense precursor ions per full MS scan were isolated in the quadrupole using a 1.6 m/z isolation window and fragmented by higher-energy collisional dissociation (HCD) using stepped normalized collision energies of 20, 40, and 60 %. Dynamic exclusion was enabled with an exclusion duration of 6 s using a mass tolerance of ±6 ppm. The automatic gain control (AGC) targets were set to 4 × 10⁵ for full MS and 5 × 10⁴ for MS/MS acquisitions.

### 2.5. LC-MS/MS data processing and metabolite annotation

The LC-MS/MS raw data files were pre-processed using Compound Discoverer 3.1 (Thermo Fisher Scientific, USA) with ESI+ and ESI-mode datasets processed separately.

Mass detection, chromatogram building, and peak deconvolution were performed to detect chromatographic features defined by unique combinations of accurate mass-to-charge ratio (m/z) and retention time (RT). Detected features were aligned across all samples and peak areas were quantified based on extracted ion chromatograms. Isotope and compound grouping were also applied.

Analytical reproducibility, retention time stability, and mass accuracy were monitored using pooled quality control (QC) samples and a QC mix of reference standards injected throughout the analytical sequence.

Putative compound annotation was performed based on accurate mass, retention time, and MS/MS fragmentation spectra. MS/MS spectra associated with detected features were matched against an in-house MS² spectral library as well as publicly available databases, both MS1 and MS/MS using Compound Discoverer. Spectral matching was performed using a mass tolerance of 5 ppm. Candidate annotations were assigned based on accurate mass agreement and MS/MS spectral similarity. When available, retention time agreement with in-house reference compounds was used to increase annotation confidence.

Compound annotations were reported according to Radi et al., in which a level-1 identification was assigned when a feature achieved ≥70 % spectral similarity to entries in the in-house mzVault library. When necessary, MS/MS spectra were manually inspected and matched against reference standards to confirm metabolite identities.

Potential metabolite exchange events were identified through automated analysis in R. Low-confidence signals were first removed using predefined filtering criteria. For extract analyses, peak intensities were normalized to the oocyte count of each replicate. Log2-transformed data were then subjected to univariate statistical analysis using a paired t-test. Volcano plots were generated using a significance threshold of p < 0.05 and an absolute log2 fold change > 0.5 to identify compounds exhibiting significant changes in abundance across time points. Manual inspection of raw data was subsequently performed to retain only high-quality peaks with clear chromatographic apices, consistent peak shapes, stable baselines, high signal-to-noise ratios, and reproducible retention times across replicates, while excluding noise-derived or poorly integrated features.

### 2.6. Molecular docking simulation

Molecular docking analysis was carried out for SLC16A10 to computationally assess the potential of different substrates binding to the transporter. The receptor structure was obtained from the Protein Data Bank (PDB; rcsb.org) [25] based on experimental determination via cryo-EM by Bågenholm et al. (2025) [26]. The structure (PDB ID: 9HHQ, 3.5Å resolution) was pre-processed using Meeko (version 0.6.1 Forli Lab, 2024) [27], RDKit (version 2023.09.6) [28] and OpenMM (version 8.2.0) [29].

The cargo-binding pocket was used to define the docking search space. The pocket is supported by two independent structural studies: Bågenholm et al. (2025) [26] identified it as a conserved inward-open cleft, with key lining residues confirmed by mutagenesis to be critical for transport activity, and Tassinari et al. (2025) [30] determined the cryo-EM structure of MCT10 bound to thyroxine (PDB ID: 9GSZ; 3.8 Å resolution), providing direct experimental validation of ligand binding at this site. The docking box grid was centered on this pocket at (0, 1, 1) with dimensions of 30×23×30 Å^3^, set to encompass the full extent of the binding cleft.

Three-dimensional structures of all ligands, including positive controls, negative controls and novel substrates were obtained from PubChem [31] and prepared using Meeko and RDKit. Positive controls consisted of known binders of SLC16A10, namely Tryptophan, Phenylalanine, L-DOPA, Thyroxine and Tyrosine. For negative controls, Histidine and Leucine were selected, which are known not to be transported by SLC16A10. Hippuric acid and 3-phenyllactic acid were docked as potential novel substrates.

Docking was performed using AutoDock Vina (version 1.2.7) [32–34] with an exhaustiveness parameter of 32 for improved sampling. For each ligand, the top 10 binding poses were retained and ranked by predicted binding affinity. Docked poses were visualized and analyzed using PyMOL (the PyMOL Molecular Graphics System, Version 3.1.3 Schrödinger, LLC), and atom-level interactions of the top scoring poses for Phenylalanine, Hippuric acid and 3-phenyllactic acid were identified with PLIP 2025 [35] for visualization.

Finally, the locations of the docked poses within the receptor were analyzed with respect to the presumed binding pocket. The afore mentioned cryo-EM structure of MCT10 bound to thyroxine (PDB ID: 9GSZ; 3.8Å resolution) [30] served as a reference for the likely binding pocket within the transporter. The MCT10 structure used in the docking analysis (PDB ID: 9HHQ, 3.5 Å resolution) [26] was aligned to this reference, yielding an overall root mean square deviation (RMSD) of 1.100 Å. Residues within 3 Å of thyroxine after alignment were defined as the pocket, and a 3D convex hull was constructed around these residues using Delaunay triangulation. For each of the top 10 docked binding poses of every substrate, the centroid was calculated; a pose whose centroid fell within the hull was considered to lie in or near the protein’s presumed canonical binding pocket.

## 3. Results

### 3.1. Blood serum as a metabolite library for *Xenopus* oocyte transport assays

A workflow for capturing metabolite exchange in transporter-expressing *Xenopus* oocytes was established in this study, enabling separate profiling of intracellular and extracellular metabolites (Figure 1). Although uptake-focused assays are commonly used in oocyte-based transporter characterization, recent work has shown that extracellular measurements of metabolite exchange between eukaryotic and bacterial cells and human serum can be detected conveniently using untargeted metabolomics [13,14]. Analyzing both the cellular extract and the surrounding serum can potentially provide a more comprehensive view of metabolite exchange and yield deeper insight into how transporter expression in oocytes reshapes both intra– and extracellular metabolite profiles.

**Figure 1.**
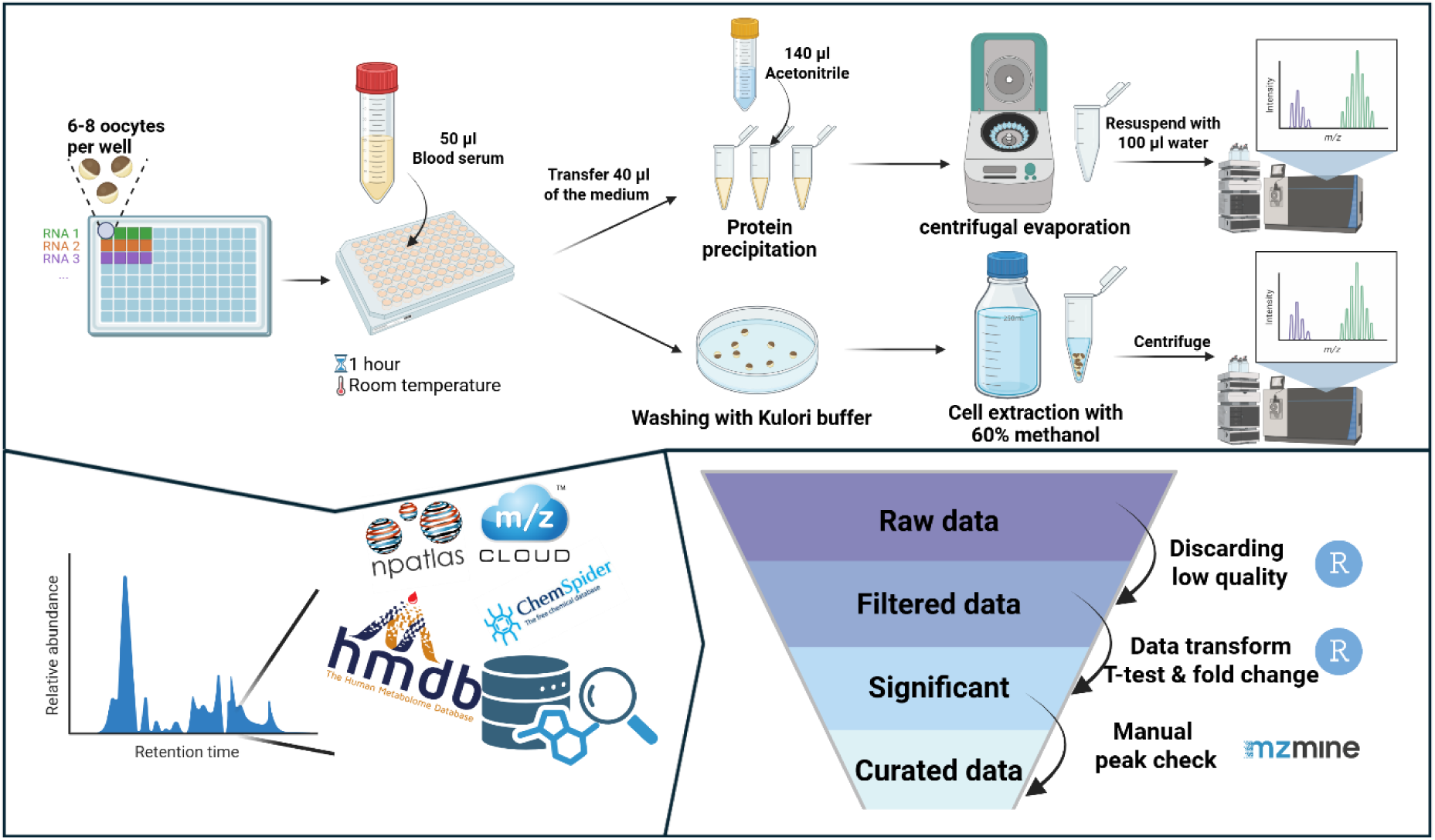
The procedure of human blood serum incubation for transporter characterization as used in this study.

As a proof-of-concept study, five SLC transporters were selected and expressed by mRNA injection in *Xenopus* oocytes. The five transporters have experimentally validated substrates covering amino acids, organic acids, bile acids, and cofactors, and they operate through distinct transport mechanisms: sodium-coupled, proton-coupled, or facilitative transport (Table 1).

**Table 1.** SLC transporters analyzed in this study.

| Name | Synonyms | Reported substrates | Mechanism of action | Reference |
| --- | --- | --- | --- | --- |
| SLC10A2 | ASBT | Bile acids | Sodium symport | [18] |
| SLC10A6 | SOAT | Dehydroepiandrosterone sulfate, estrone-3-sulfate, pregnenolone sulfate, tauroolithocholic acid-3-sulfate | Sodium symport | [19] |
| SLC13A2 | NaDC-1 | Succinic acid, citric acid | Sodium symport | [20] |
| SLC16A10 | TAT1 | Tryptophan, phenylalanine, tyrosine, L-DOPA, thyroid hormones | Facilitated diffusion | [21] |
| SLC46A1 | HCP1 | Folate | Proton symport | [22] |

Uptake assays were performed for each transporter, using GFP-injected oocytes lacking the transporter-encoding mRNA as a negative control, and metabolite exchange was monitored by LC-MS. MS/MS acquisition was conducted exclusively on QC samples, and data processing was carried out using Compound Discoverer. A total of 6027 and 2845 features were detected in ESI+ and ESI– modes, respectively (Table 2).

**Table 2.** Overview of LC-MS/MS results of serum QC samples obtained following preprocessing using Compound Discoverer 3.1.

|  | ESI+ | % All Compounds | ESI- | % All Compounds |
| --- | --- | --- | --- | --- |
| Features | 6027 |  | 2845 |  |
| MS2 | 1445 | 24 % | 717 | 25 % |
| No MS2 | 4582 | 76 % | 2128 | 75 % |
| MS2 Preferred Ion | 1008 | 17 % | 708 | 25 % |
| MS2 Other Ion | 437 | 7 % | 9 | 0.3 % |
| Level 1 (In-house database, MzVault >70 %) | 118 | 2 % | 71 | 2 % |
| Level 2 (Probable structure >70 %) | 49 | 0.8 % | 11 | 0.4 % |
| Level 3 (Tentative Candidate, match to predicted composition) | 4303 | 71 % | 1980 | 70 % |
| Level 4 (Molecular formula) | 495 | 8 % | 133 | 5 % |
| Level 5 (Mass) | 1062 | 18 % | 650 | 23 % |

We first examined metabolite exchange between *Xenopus* oocytes and human blood serum using principal components analysis (PCA). In the PCA of positive-mode LC-MS data, samples of native serum and serum incubated with GFP-expressing oocytes separated along PC2, indicating that the presence of oocytes induces a detectable shift in the serum metabolite profile (Figure 2). In contrast, ESI-mode data did not discriminate the two conditions (Figure S1). PCA analysis of the SLC transporter-expressing groups also did not reveal transporter-specific clustering, suggesting that introducing individual SLCs does not *per se* lead to a strong global metabolic signature in the oocyte (data not shown). Although previous studies using *E. coli* or mammalian cell lines have reported more pronounced metabolite profile changes following serum incubation, the comparatively subtle shifts observed in *Xenopus* oocytes may reflect their relatively clean endogenous metabolic background, potentially due to limited membrane-associated metabolic activity. Furthermore, the minor differences in the PCA analysis reflect the near-dormant background transport activity of the *Xenopus* oocytes in respect to the human blood serum metabolites.

**Figure 2.**
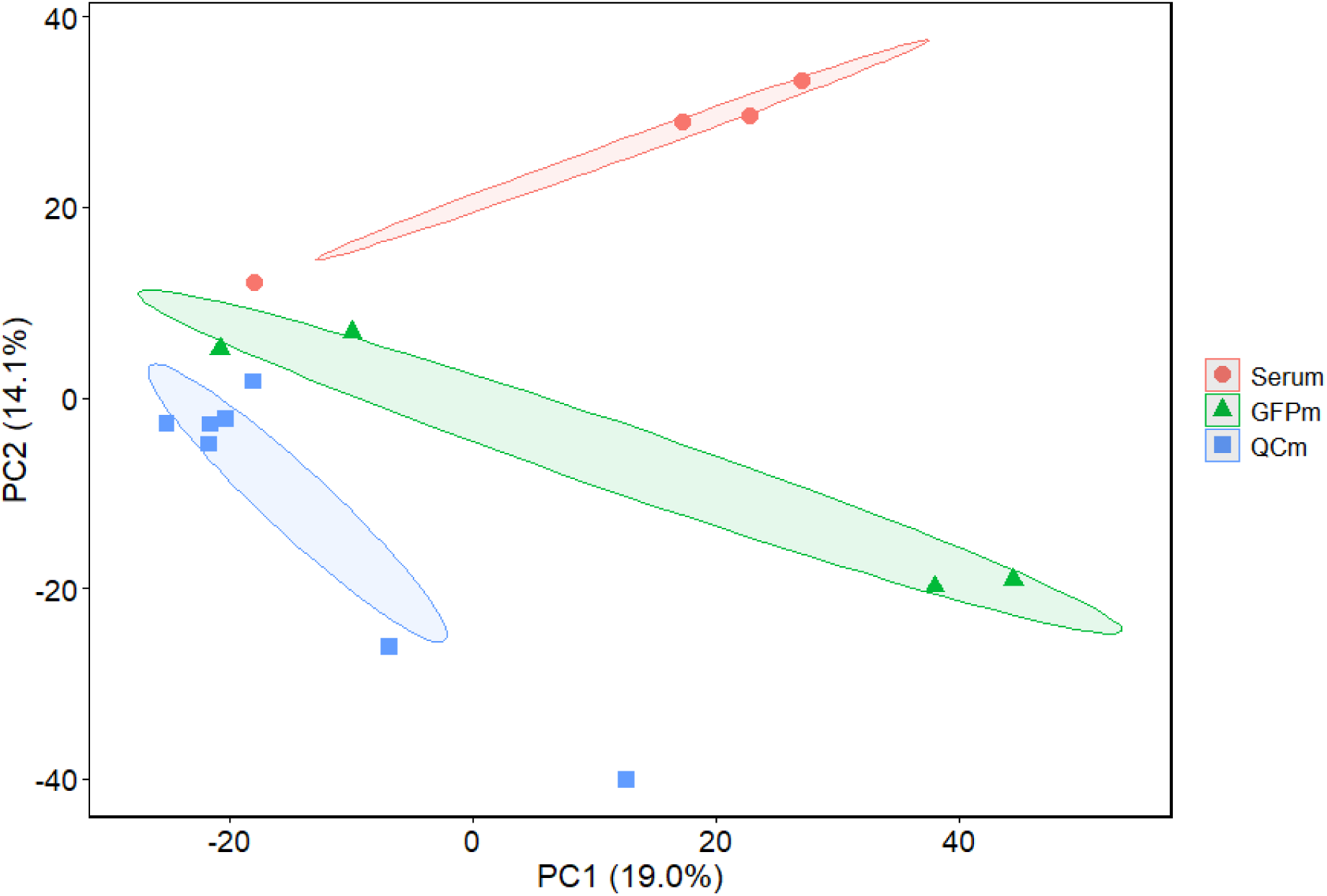
Principal component analysis (PCA) of serum, serum incubated with GFP mRNA-injected *Xenopus* oocytes, and QC samples based on positive-ion (ESI+) mode LC-MS/MS data.

### 3.2. Metabolite exchange profiling of *Xenopus* oocytes expressing SLC transporters

By comparing the metabolic footprints of *Xenopus* oocytes expressing SLC transporters with those expressing GFP alone as controls, differential metabolites were identified. In intracellular extracts, increased metabolite abundance was interpreted as potential import, whereas decreased abundance was considered indicative of efflux. For the incubated serum, the expected trends were reversed, with decreases suggesting uptake by the oocyte and increases potentially reflecting transporter-mediated secretion (Figure 3, Figure S2).

**Figure 3.**
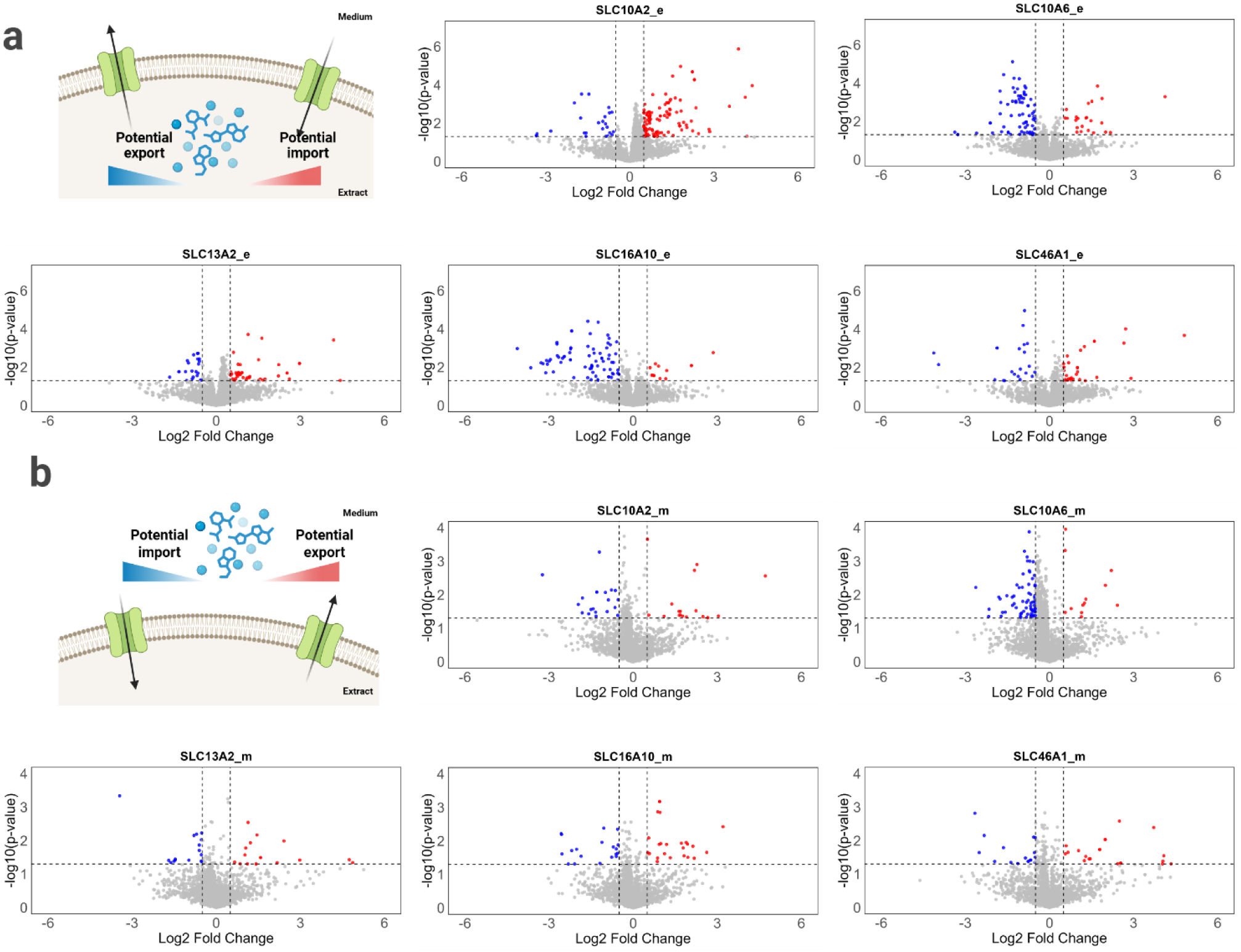
Metabolome volcano plots of intracellular (a) and extracellular (b) LC–MS/MS features acquired in ESI+ mode. Red and blue dots indicate features significantly enriched or depleted, respectively, in transporter-expressing oocytes relative to controls; grey dots indicate non-significant features. Significance was defined as p < 0.05 and absolute log₂ fold change > 0.5, indicated by the horizontal and vertical dashed lines, respectively. Each data point represents the average of four independent experiments, with each replicate consisting of 6–8 oocytes

For each transporter, volcano plot analysis revealed signatures consistent with both import and efflux events. While symporters such as SLC10A2, SLC10A6, SLC13A2, and SLC46A1 are classically associated with ion-gradient-driven uptake, SLC16A10 functions as a facilitated-diffusion transporter, inherently supporting bidirectional transport driven by substrate concentration gradients, which is consistent with the observation of both uptake and release signatures.

Because automated annotation can introduce false positives, to quantify these transport-associated changes with higher confidence, manual inspection of raw chromatographic peaks was performed to remove features with poor peak shape or low signal quality (Figure S3). The resulting curated dataset, therefore, represents a condensed set of higher-confidence exchange events (Table 3, Table S4). Across transporters, a greater number of exchange events were detected in oocyte extracts than in the serum medium in both ionization modes. This might indicate a higher sensitivity for intracellular measurements under the current workflow or, more likely, simply the fact that serum contained many more metabolites than did the cells and that antiporter activity was predominant. Consistent with this trend, fewer metabolite exchange events in the extract were detected in oocytes expressing SLC46A1, a proton-coupled folate symporter, compared with the other transporters examined. This observation is in line with the relatively small proton gradient between human blood serum and the *Xenopus* oocyte cytosol, which is expected to be less pronounced than the sodium gradient that drives transport for the sodium-coupled symporters analyzed in this study.

**Table 3.** Statistics of potential transport events identified by LC-MS/MS. The total number of candidate transport events and the subsets retained after manual peak inspection (curated) are shown. A detailed list of potential transport events is provided in Supplementary Table S4

| Transporter | ESI+ |  |  |  | ESI- |  |  |  |
| --- | --- | --- | --- | --- | --- | --- | --- | --- |
|  | Medium |  | Extract |  | Medium |  | Extract |  |
|  | Features | Curated | Features | Curated | Features | Curated | Features | Curated |
| SLC10A2 | 38 | 9 | 127 | 38 | 24 | 2 | 175 | 24 |
| SLC10A6 | 80 | 20 | 90 | 17 | 127 | 44 | 100 | 10 |
| SLC13A2 | 31 | 5 | 62 | 10 | 23 | 3 | 58 | 15 |
| SLC16A10 | 42 | 11 | 80 | 16 | 51 | 17 | 66 | 15 |
| SLC46A1 | 36 | 7 | 49 | 6 | 77 | 20 | 39 | 3 |

### 3.3. Identification of substrates for SLC transporters

After identifying potential metabolite transport events, we sought to contextualize our findings with existing knowledge by examining whether the detected metabolites correspond to substrates previously reported to be transported by the SLC transporters.

SLC10A2 is a Na⁺-coupled bile acid symporter that mediates bile acid transport driven by the transmembrane sodium gradient. It has been reported to transport a broad range of bile acids. Given the higher sodium concentration in human blood serum (135-146 mM [36]) compared with the intracellular sodium concentration of *Xenopus* oocytes (4-23 mM [37]), the gradient is theoretically favorable to facilitate sodium-coupled import. Consistent with this, a potential uptake event for glycocholic acid was detected in intracellular extracts of oocytes expressing SLC10A2. The identity of glycocholic acid was further confirmed by matching the mass spectral features to in-house LC-MS reference data (Figure S4). While several other bile acids were detected, their relatively low peak intensities indicate less efficient detection and/or lower apparent accumulation under the experimental conditions.

SLC16A10, also known as MCT10, is a member of the monocarboxylate transporter family that is distinct from other family members in its ability to facilitate the diffusion of aromatic amino acids, and, as reported later, thyroid hormones. Both uptake and efflux have been reported for SLC16A10, consistent with a bidirectional, gradient-driven mode of transport.

Consistent with existing knowledge, LC-MS/MS analysis of SLC16A10-expressing *Xenopus* oocytes acquired in ESI+ mode revealed potential transport of the aromatic amino acids tryptophan and phenylalanine (Figure S5, S6). Compared with GFP-expressing controls, both metabolites exhibited significantly reduced intracellular peak intensities, consistent with transporter-mediated efflux (Figure 5). For tyrosine, a feature corresponding to its in-source fragment generated by neutral loss of NH₃ was detected in ESI+ mode and showed a pattern indicative of secretion. Alignment of retention time and MS/MS spectra confirmed this feature as fragmented tyrosine (Figure S7). In addition, ESI-mode analysis also supported an export event for tryptophan. Together, these observations support the successful identification of the three canonical substrates of SLC16A10 in this system. L-DOPA, another reported substrate of SLC16A10, was not captured as a level-1 identification in the LC-MS/MS dataset.

**Figure 4.**
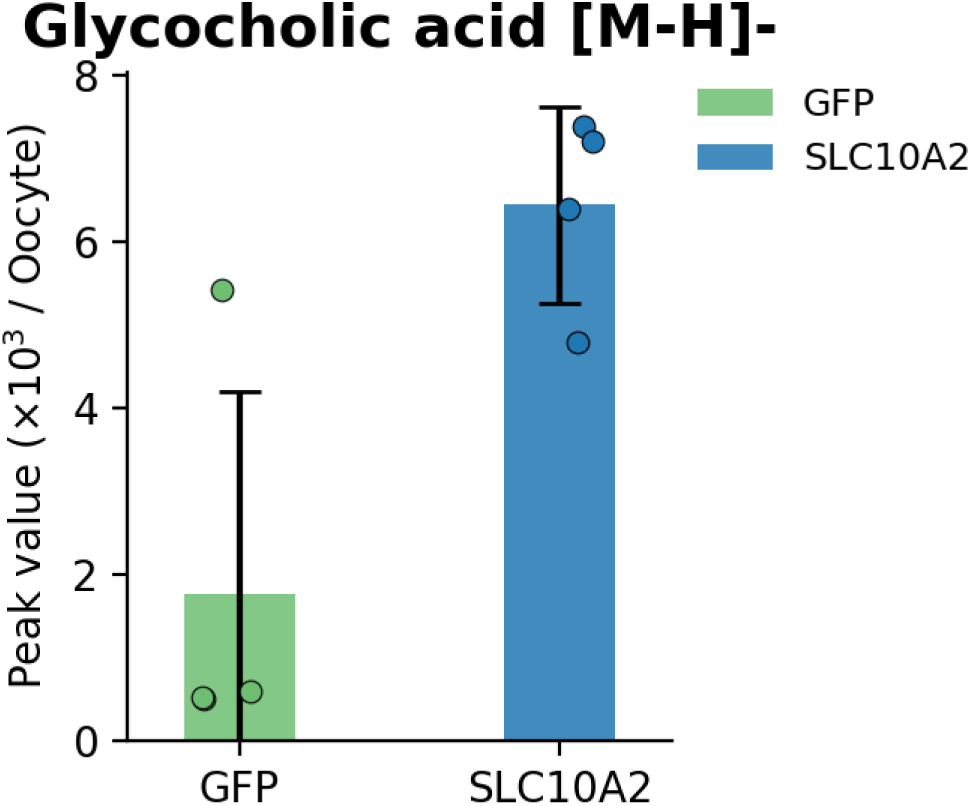
Bar plot showing glycocholic acid peak intensity in oocyte extract samples from SLC10A2-expressing and GFP-control oocytes. Data represent the average of four independent experiments, with each replicate consisting of 6-8 oocytes

**Figure 5.**
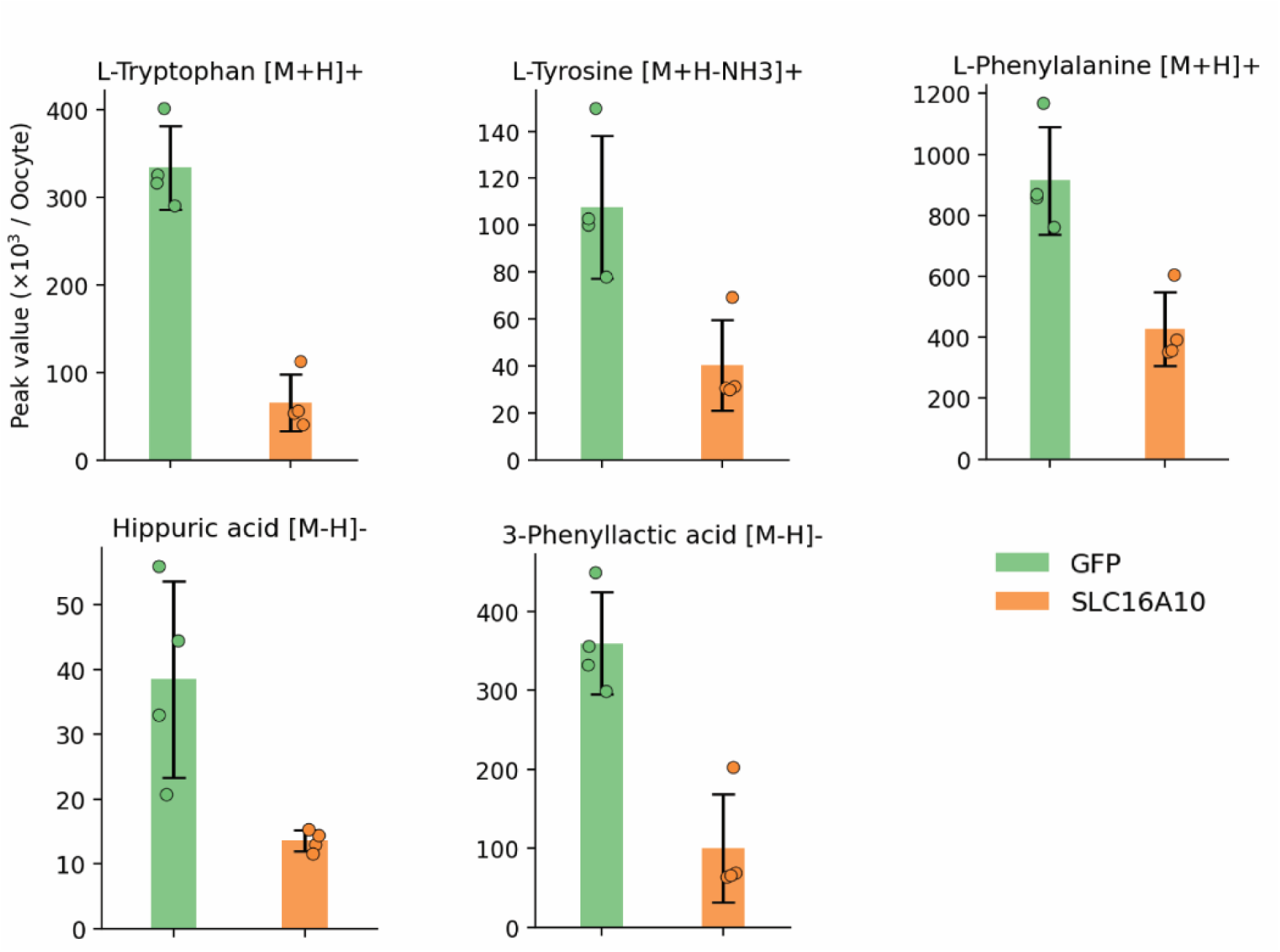
Intracellular peak intensities of five level-1 identified metabolites showing significant differences between SLC16A10-expressing *Xenopus* oocytes and GFP-expressing controls. Data represent the average of four independent experiments, with each replicate consisting of 6-8 oocytes

The successful identification of SLC16A10’s known substrates from both ESI+ and ESI-modes provides a solid foundation and confidence for discovering additional potential substrates within the dataset. Among the annotated and manually curated features, hippuric acid and 3-phenyllactic acid were identified and confirmed by matching the signals to the in-house MS/MS reference library, revealing confirmation of the compound (Figure S8, S9).

### 3.4. Molecular docking of reported and new potential substrates of SLC16A10

The binding affinity of substrates within the transporter’s substrate binding pocket is prerequisite to transporter selectivity [38,39]. To investigate the interactions between SLC16A10 and small molecule substrates, we performed a series of molecular docking analyses using its known substrates, two validated non-substrates (histidine and leucine [40]), and the two potential substrates identified in this study. Consistent with the expected selectivity pattern, SLC16A10 exhibited markedly different binding affinities toward substrates versus non-substrates, as reflected by the predicted free binding energy distribution (Figure 6, Table S5). The binding affinity profile of hippuric acid and 3-phenyllactic acid showed a similar distribution to original substrates of SLC16A10, suggesting that overall structural similarity, rather than the presence of an amino acid functional group, may represent a key determinant of substrate recognition by SLC16A10.

**Figure 6.**
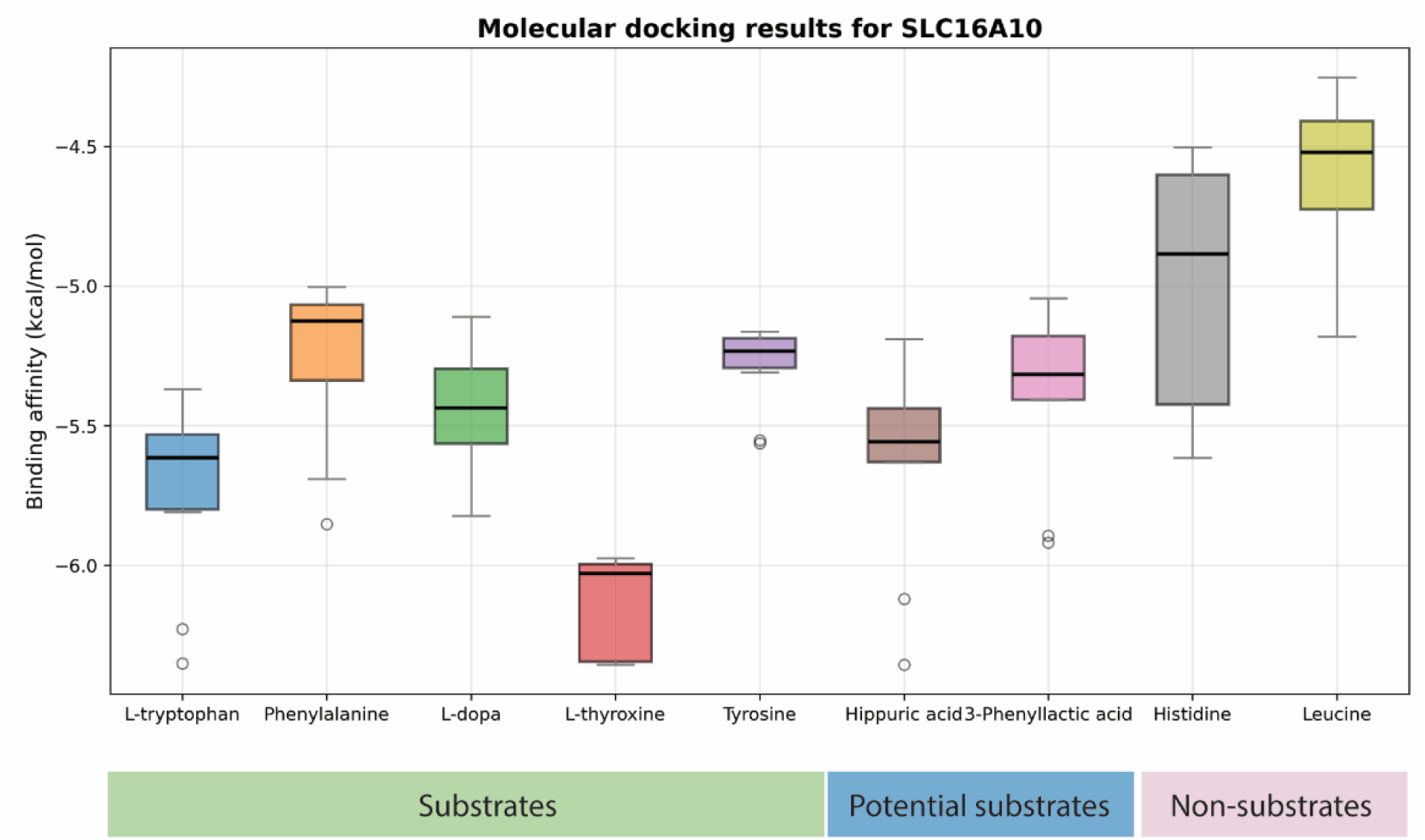
Predicted binding free energies of selected SLC16A10 ligands from the top 10 AutoDock Vina poses with the strongest binding affinities, grouped as validated substrates, potential substrates, or non-substrates. Lower Gibbs free energy values indicate stronger predicted binding.

We further examined the binding locations of the highest-affinity predicted poses, using the cryo-EM structure of MCT10 bound to thyroxine as a reference for the likely canonical binding pocket. For the known substrates, most (70-100%, and ≥80% for all but one substrate) of the top 10 predicted poses fell within this pocket, defined as the residues within 3 Å of the bound ligand in the reference structure (Figure S10-S11). For the potential new substrates, hippuric acid and 3-phenyllactic acid, AutoDock Vina placed 80% of highest-affinity poses in the same pocket. Conversely, for the non-substrates, most (70%) of the top 10 poses were located outside the presumed canonical binding pocket. Altogether, the predicted poses fell within the canonical binding pocket for known and candidate substrates but not for non-substrates, discriminating substrates from non-substrates and supporting hippuric acid and 3-phenyllactic acid as plausible substrates.

## 4. Discussion

Identifying (often multiple or promiscuous) substrates of transporter activity remains a major challenge in transporter characterization. Previous studies, including our own, have demonstrated that compound libraries combined with LC-MS analysis can be powerful tools for transporter profiling, enabling sensitive and quantitative detection of transport activity toward defined xenobiotics and endogenous metabolites, albeit they cannot discriminate pleiotropic effects. However, the predefined composition and limited chemical diversity of such libraries inherently constrain their ability to uncover non-intuitive or unexpected substrates. By contrast, human blood serum contains thousands of endogenous metabolites spanning a broad physicochemical space (e.g. [41]), making it an attractive and biologically relevant alternative metabolite library for unbiased substrate discovery.

In this study, we established a *Xenopus* oocyte-based transporter characterization platform aimed at exploring transporter substrate ranges in an unbiased manner. By incubating oocytes in human blood serum and profiling intracellular and extracellular metabolite pools, this workflow enables the simultaneous detection of potential uptake and efflux events, providing information for the identification of transporter-associated metabolite exchanges.

Matching detected metabolite exchange events with previously reported substrates confirmed the successful identification of known substrates for both SLC10A2 and SLC16A10. For both transporters, the matched signals were predominantly identified in intracellular extracts of oocytes rather than in the incubation medium, suggesting that extract-based analysis, when combined with the current LC-MS/MS workflow, provides higher sensitivity for capturing at least some transporter-mediated exchange events and yields greater confidence than medium analysis.

Importantly, features exhibiting significant abundance changes were subjected to manual inspection of the raw LC-MS/MS data to evaluate peak shape, signal-to-noise ratio, and chromatographic consistency. This additional quality-control step reduced the likelihood of false-positive annotations arising from noise, co-eluting interferences, or poorly resolved features, thereby enhancing the robustness of the curated dataset (Figure S3). Nevertheless, it is important to note that even after manual inspection, the alignment of MS/MS spectra to reference standards remains essential when interpreting the dataset, even after automated filtering and manual data-cleaning procedures (Figure S4-9). In particular, level-3 identifications offer only tentative structural information, as they are primarily based on accurate mass and predicted molecular formulae. Additional MS/MS evidence is therefore required to increase confidence in metabolite assignment. Furthermore, most observed mass spectra are not represented in standard metabolite libraries, highlighting the potential need for de novo identification strategies [42].

SLC16A10 is known to facilitate the diffusion of aromatic amino acids and thyroid hormones. In this study, significant decreases in intracellular tryptophan, phenylalanine, and tyrosine were observed in SLC16A10-expressing oocytes compared with GFP controls, whereas other reported substrates were not detected under the applied analytical conditions. The reduced intracellular abundance of aromatic amino acids is consistent with transporter-mediated efflux and can be rationalized by two factors: First, the total free amino acid concentration in *Xenopus* oocytes (∼16.4 mmol L⁻¹) [43] is substantially higher than that in human serum (∼3.3 mmol L⁻¹) [44], establishing a concentration gradient that favors efflux. Second, transporter expression is initiated during the three-day incubation period following mRNA injection, during which oocytes are maintained in amino-acid-free Kulori buffer. Gradient-driven efflux during this phase may reduce intracellular baseline levels prior to serum incubation, thereby influencing the measured intracellular metabolite abundances.

In addition to aromatic amino acids, two aromatic monocarboxylates, hippuric acid and 3-Phenyllactic acid, were identified as transported by SLC16A10. Their identities were confirmed by matching LC-MS/MS spectra to in-house reference standards, achieving level 1 identification. Although these compounds are not canonical substrates of SLC16A10, their structural features share similarities with aromatic amino acids, particularly the presence of an aromatic ring and a carboxylate group. This observation suggests that substrate recognition by SLC16A10 may be influenced more broadly by aromaticity and carboxylate functionality rather than strict amino-acid identity. Molecular docking analyses supported this interpretation, showing higher predicted binding affinities for hippuric acid and 3-phenyllactic acid compared with validated non-substrate amino acids (Figure 6).

We also successfully identified a potential import event for glycocholic acid, a reported substrate of SLC10A2, the Na⁺-bile acid symporter. While several other bile acids were detected in the samples, their relatively low signal intensities limited confident interpretation, likely reflecting suboptimal detection sensitivity under the applied analytical conditions or a low content of bile acid presence in the blood serum.

It is also noteworthy that under the current analytical workflow, several expected substrates were not captured. For example, citrate and succinate, which are anticipated substrates of SLC13A2, were not observed as differential features, and folate was not detected in the LC-MS/MS dataset. Extraction efficiency and analytical sensitivity are not uniform across metabolite classes, and LC-MS/MS acquisition parameters can bias the metabolites recovered and detected. Accordingly, these absences are likely attributable to methodological limitations rather than lack of transport activity. In particular, small organic acids such as citrate and succinate typically exhibit poor chromatographic retention and reduced sensitivity under generic reversed-phase LC conditions, while folate and related vitamins often require tailored extraction procedures, chromatographic separation, and ionization settings for reliable detection [45–47].

Across all transporters examined, signatures consistent with bidirectional metabolite exchange were observed. While the bidirectional transport signals detected for SLC16A10 are consistent with its facilitated-diffusion mechanism, symporters are classically associated with gradient-driven substrate uptake. Nevertheless, apparent efflux events observed for symporters may arise from local ion micro-gradients at the oocyte membrane or from secondary metabolic effects induced by transporter expression. These counterintuitive observations warrant more careful consideration and mechanistic interpretation.

In summary, this study introduces a generalizable framework for transporter substrate discovery by leveraging biological complexity rather than predefined compound sets. By decoupling substrate discovery from prior assumptions and enabling systematic refinement through targeted methodological optimization, this platform provides a promising foundation for expanding transporter annotation and for probing transporter promiscuity across diverse chemical spaces.

## Author Contributions

Conceptualization: LDS, CGAR, DBK, IB. Methodology: YZ, LDS, DR, FCS, ASD. Investigation: YZ, LDS, FCS, ASD. Formal analysis: YZ, FCS. Data curation: YZ, DR. Visualization: YZ, FCS. Funding acquisition: IB, CGAR. Writing and editing: YZ, LDS, DR, FCS, IB, DBK, CGAR.

## Acknowledgements

This project has received funding from The European Research Council (ERC) under the European Union’s Horizon 2020 research and innovation programme (YEAST-TRANS, grant agreement No. 757384) and the Novo Nordisk Foundation (Grant agreement No. NNF20OC0060809 and NNF14CC0001 and NNF24SA0100980).

