## Supplementary materials for "Unlocking substrate specificities of human solute carrier proteins using untargeted metabolomics"

**Supplementary Table 1.** Oligonucleotide primers used in this study.

| **Name** | **Sequence (5’ to 3’)** | **Orientation** | **Description** |
|---|---|---|---|
| PR-36701 (SLC10A2_frog_F) | GGCTTAAUATGAATGATCCAAATTCTTGTGTGGATAA | → | Amplification of SLC10A2 to be cloned into pCfB5245 |
| PR-36702 (SLC10A2_frog_R) | GGTTTAAUTTATTTTTCATCTGGCTGAAATCCTCCA | ← | |
| PR-36703 (SLC10A6_frog_F) | GGCTTAAUATGAGAGCTAATTGTTCTTCTTCTTCTGCTT | → | Amplification of SLC10A6 to be cloned into pCfB5245 |
| PR-36704 (SLC10A6_frog_R) | GGTTTAAUTTATTCACAAGATGTAATGTGTCCCACTGGTT | ← | |
| PR-36705 (SLC13A2_frog_F) | GGCTTAAUATGGCAACTTGCTGGCAAGC | → | Amplification of SLC13A2 to be cloned into pCfB5245 |
| PR-36706 (SLC13A2_frog_R) | GGTTTAAUCTACGGGGAAGGCGTGGT | ← | |
| PR-36707 (SLC16A10_frog_F) | GGCTTAAUATGGTGCTGTCTCAGGAAGAACC | → | Amplification of SLC16A10 to be cloned into pCfB5245 |
| PR-36708 (SLC16A10_frog_R) | GGTTTAAUTTAAATAATAGAATCAGATTCTTTTTTAAACATTCCAGAAGAA | ← | |
| PR-36709 (SLC46A1_frog_F) | GGCTTAAUATGGAAGGAAGTGCAAGTCCG | → | Amplification of SLC46A1 to be cloned into pCfB5245 |
| PR-36710 (SLC46A1_frog_R) | GGTTTAAUTCAGGGTGATTGAGGGAATTGTTGG | ← | |

**Supplementary Table 2.** Plasmids used in this study.

| Name | Description | Reference |
|---|---|---|
| pCfB5245 | USER-compatible *Xenopus* expression vector | [19] |
| pCfB13607(p5245_SLC10A2_oocyte) | Oocyte mRNA synthesis plasmid for expression of SLC human transporter SLC10A2 | This work |
| pCfB13608(p5245_SLC10A6_oocyte) | Oocyte mRNA synthesis plasmid for expression of SLC human transporter SLC10A6 | This work |
| pCfB13609(p5245_SLC13A2_oocyte) | Oocyte mRNA synthesis plasmid for expression of SLC human transporter SLC13A2 | This work |
| pCfB13610(p5245_SLC16A10_oocyte) | Oocyte mRNA synthesis plasmid for expression of SLC human transporter SLC16A10 | This work |
| pCfB13611(p5245_SLC46A1_oocyte) | Oocyte mRNA synthesis plasmid for expression of SLC human transporter SLC46A1 | This work |

**Supplementary Table 3.** Synthesized codon optimized genes of SLC transporters.

| Name | Construct length (Insert + Adapters) | Construct sequence |
|---|---|---|
| SLC10A1 | 1094 | CAATCCGCCCTCACTACAACCGATGGAAGCTCACAATGC<br>TTCTGCTCCATTTAATTTACACTGCCACCAAATTTTGGA<br>AAAAGACCAACAGATCTGGCTCTGTCTGTGATTCTGGTG<br>TTTATGCTGTTTTTTATTATGCTGTCTCTGGGATGTACAAT<br>GGAATTTTCTAAAATTAAAGCTCACCTGTGGAAACCAAA<br>AGGACTGGCTATTGCTCTGGTGGCTCAGTATGGAATTAT<br>GCCACTGACAGCTTTTGTGCTGGGAAAAGTGTTTAGAC<br>TGAAAAATATTGAAGCTCTGGCTATTCTGGTGTGTGGAT<br>GTTCTCCAGGAGGAAATCTGTCTAATGTGTTTTCTCTGG<br>CTATGAAAGGAGATATGAATCTGTCTATTGTGATGACAA<br>CATGTTCTACATTTTGTGCTCTGGGAATGATGCCACTGCT<br>GCTGTATATTTATTCTAGAGGAATTTATGATGGAGATCTG<br>AAAGATAAAGTGCCATATAAAGGAATTGTGATTTCTCTG<br>GTGCTGGTGCTGATTCCATGTACAATTGGAATTGTGCTG<br>AAATCTAAAAGACCACAGTATATGAGATATGTGATTAAA<br>GGAGGAATGATTATTATTCTGCTGTGTTCTGTGGCTGTGA<br>CAGTGCTGTCTGCTATTAATGTGGGAAAATCTATTATGTT<br>TGCTATGACACCACTGCTGATTGCTACATCTTCTCTGATG<br>CCATTTATTGGATTTCTGCTGGGATATGTGCTGTCTGCTC<br>TGTTTTGTCTGAATGGAAGATGTAGAAGAACAGTGTCTA<br>TGGAAACAGGATGTCAGAATGTGCAGCTGTGTTCTACA<br>ATTCTGAATGTGGCTTTTCCACCAGAAGTGATTGGACCA<br>CTGTTTTTTTTTCCACTGCTGTATATGATTTTTCAGCTGG<br>GAGAAGGACTGCTGCTGATTGCTATTTTTTGGTGTTATG<br>AAAAATTTAAAACACCAAAAGATAAAACAAAAATGATT<br>TATACAGCTGCTACAACAGAAGAAACAATTCCAGGAGC<br>TCTGGGAAATGGAACATATAAAGGAGAAGATTGTTCTCC<br>ATGTACAGCTTAACTACTCTGGCGTCGATGAGGGA |
| SLC10A2 | 1091 | CAATCCGCCCTCACTACAACCGATGAATGATCCAAATTC<br>TTGTGTGGATAATGCTACAGTGTGTTCTGGAGCTTCTTGT<br>GTGGTGCCAGAATCTAATTTTAATAATATTCTGTCTGTGG<br>TGCTGTCTACAGTGCTGACAATTCTGCTGGCTCTGGTGA<br>TGTTTTCTATGGGATGTAATGTGGAAATTAAAAAATTTCT<br>GGGACACATTAAAAGACCATGGGGAATTTGTGTGGGATT<br>TCTGTGTCAGTTTGGAATTATGCCACTGACAGGATTTATT<br>CTGTCTGTGGCTTTTGATATTCTGCCACTGCAGGCTGTG<br>GTGGTGCTGATTATTGGATGTTGTCCAGGAGGAACAGCT<br>TCTAATATTCTGGCTTATTGGGTGGATGGAGATATGGATC<br>TGTCTGTGTCTATGACAACATGTTCTACACTGCTGGCTCT<br>GGGAATGATGCCACTGTGTCTGCTGATTTATACAAAAAT<br>GTGGGTGGATTCTGGATCTATTGTGATTCCATATGATAAT<br>ATTGGAACATCTCTGGTGTCTCTGGTGGTGCCAGTGTCT<br>ATTGGAATGTTTGTGAATCACAAATGGCCACAGAAAGCT<br>AAAATTATTCTGAAAATTGGATCTATTGCTGGAGCTATTC |

| | | |
|---|---|---|
| | | TGATTGTGCTGATTGCTGTGGTGGGAGGAATTCTGTATCAGTCTGCTTGGATTATTGCTCCAAAACTGTGGATTATTGGAACAATTTTTCCAGTGGCTGGATATTCTCTGGGATTTCTGCTGGCTAGAATTGCTGGACTGCCATGGTATAGATGTAGAACAGTGGCTTTTGAAACAGGAATGCAGAATACACAGCTGTGTTCTACAATTGTGCAGCTGTCTTTTACACCAGAAGAACTGAATGTGGTGTTTACATTTCCACTGATTTATTCTATTTTTCAGCTGGCTTTTGCTGCTATTTTTCTGGGATTTTATGTGGCTTATAAAAAATGTCACGGAAAAAATAAAGCTGAAATTCCAGAATCTAAAGAAAATGGAACAGAACCAGAATCTTCTTTTTATAAAGCTAATGGAGGATTTCAGCCAGATGAAAAATAACTACTCTGGCGTCGATGAGGGA |
| SLC13A2 | 1823 | CAATCCGCCCTCACTACAACCGATGGCAACTTGCTGGCAAGCCTTGTGGGCCTATCGAAGTTATTTAATAGTCTTCTTCGTCCCCATCCTTCTCTTACCCCTGCCTATCTTGGTACCTTCAAAAGAGGCATACTGCGCCTACGCAATAATACTTATGGCCCTCTTCTGGTGCACCGAGGCACTTCCGCTAGCCGTTACCGCCTTGTTCCCTTTAATCCTCTTCCCCATGATGGGCATCGTTGACGCCTCTGAGGTAGCAGTTGAGTACTTGAAGGACAGCAACCTACTCTTCTTCGGCGGCTTACTTGTAGCCATAGCAGTTGAGCATTGGAACTTGCATAAGAGGATCGCACTTAGGGTCCTTCTAATCGTCGGGGTTCGACCTGCACCTCTAATCCTTGGCTTCATGCTCGTTACTGCCTTCTTGAGCATGTGGATCTCAAACACGGCAACCAGTGCAATGATGGTTCCTATCGCACATGCCGTATTAGACCAACTTCATTCAAGCCAAGCCAGTAGCAACGTCGAAGAGGGCTCAAACAACCCTACGTTCGAGCTTCAAGAGCCGAGCCCCCAAAAGGAAGTCACTAAGTTAGACAACGGGCAAGCACTCCCCGTCACTAGTGCCTCAAGCGAGGGAAGGGCACATTTATCCCAAAAGCATCTTCATTTGACTCAATGCATGAGTCTCTGCGTATGCTACAGTGCCTCAATAGGTGGGATCGCAACCCTTACTGGTACTGCCCCTAACCTCGTCCTACAAGGCCAAATCAACAGTTTGTTCCCTCAAAACGGGAACGTCGTTAACTTCGCCAGTTGGTTCAGTTTCGCATTCCCTACCATGGTCATCCTTCTATTGCTTGCATGGTTGTGGTTGCAAATATTGTTCTTAGGCTTCAACTTCCGAAAGAACTTCGGGATCGGCGAGAAGATGCAAGAGCAACAACAAGCAGCCTACTGCGTAATCCAAACGGAGCATCGCCTCTTGGGGCCCATGACTTTCGCCGAGAAGGCCATCTCCATCCTCTTCGTCATCCTTGTTTTGCTTTGGTTCACTAGGGAGCCTGGTTTCTTCCTTGGCTGGGGTAACTTGGCATTCCCGAACGCCAAAGGGGAGTCCATGGTCAGCGACGGGACTGTAGCCATCTTCATAGGTATCATCATGTTCATCATCCCTTCCAAGTTCCCTGGTTTGACTCAAGACCCTGAGAACCCTGGTAAGCTTAAGGCACCCCTTGGGTTGTTAGACTGGAAGACTGTCAACCAAAAGATGCCCTGGAACATAGTATTGCTATTGGGCGGTGGCTACGCCCTTGCCAAGGGGAGTGAGCGATCAGGCTTATCAGAGTGGCTTGGGAACAAGCTAACGCCGCTTCAAAGTGTCCCTGCACCCGCAATAGCCATAATACTTAGCTTGTTGGTCGCCACCTTCACTGAGTG |

| | | |
|---|---|---|
| | | CACCTCAAACGTAGCCACGACCACTATCTTCTTACCCATCCTCGCATCCATGGCCCAAGCAATCTGCCTTCATCCTTTATACGTTATGCTACCGTGCACTCTCGCCACTAGTCTCGCATTCATGTTACCTGTAGCCACTCCGCCCAACGCCATCGTTTTCAGTTTCGGCGACCTCAAGGTCCTAGACATGGCAAGGGCAGGCTTCCTTTTAAACATCATCGGTGTATTGATCATAGCCTTAGCAATCAACAGCTGGGGCATACCTCTATTCTCCTTGCATTCCTTCCCTTCCTGGGCCCAATCCAACACTACTGCCCAATGCCTACCCTCACTAGCAAACACCACCACGCCTTCCCCGTAGCTACTCTGGCGTCGATGAGGGA |
| SLC46A1 | 1424 | CAATCCGCCCTCACTACAACCGATGGAAGGAAGTGCAAGTCCGCCAGAGAAGCCTCGAGCACGCCCAGCAGCCGCAGTTTTGTGCCGGGGTCCTGTCGAGCCGCTCGTTTTCTTGGCAAACTTCGCCTTGGTCTTACAAGGGCCCTTGACCACCCAATACCTTTGGCATCGGTTCTCCGCCGACTTGGGGTACAACGGGACCCGGCAACGTGGTGGCTGCTCAAACCGCTCAGCAGACCCTACCATGCAAGAGGTTGAGACTCTAACTTCCCATTGGACCCTTTACATGAACGTCGGCGGTTTCTTGGTTGGTCTTTTCAGCAGTACCTTGCTAGGTGCCTGGTCCGACTCCGTAGGTAGGCGTCCTCTCTTGGTTTTGGCATCACTCGGTTTATTACTTCAAGCCCTCGTATCCGTCTTCGTCGTACAACTCCAACTTCATGTCGGTTACTTCGTTTTAGGTCGGATCCTTTGCGCATTGCTTGGGGACTTCGGTGGACTCCTAGCAGCCTCATTCGCCAGCGTAGCCGACGTCAGCAGTTCAAGGTCCCGAACGTTCCGGATGGCCTTGTTAGAGGCAAGCATAGGGGTTGCCGGCATGTTAGCAAGTCTTCTCGGCGGTCATTGGTTACGGGCCCAAGGCTACGCAAACCCTTTCTGGTTGGCATTAGCCCTACTCATCGCAATGACCCTATACGCAGCCTTCTGCTTCGGCGAGACCCTCAAGGAGCCTAAGAGTACCAGGCTTTTCACGTTCCGACATCATAGGAGTATCGTTCAACTTTACGTCGCCCCTGCCCCGGAGAAGTCACGTAAGCATCTAGCCCTCTACAGCCTTGCCATCTTCGTTGTCATCACCGTTCATTTCGGCGCACAAGACATCCTTACCCTCTACGAGCTCAGTACGCCTTTATGCTGGGACAGCAAGTTAATCGGCTACGGGAGCGCAGCACAACATTTGCCGTACTTAACTAGTCTTCTTGCCTTGAAGCTATTACAATACTGCCTTGCAGACGCCTGGGTTGCAGAGATAGGGTTGGCATTCAACATCCTTGGTATGGTTGTATTCGCCTTCGCAACCATCACTCCCTTGATGTTCACTGGTTACGGCCTATTGTTCTTGTCCCTTGTTATCACGCCTGTCATACGCGCCAAGTTGAGCAAGCTTGTTCGAGAGACTGAGCAAGGCGCACTCTTCAGTGCCGTAGCATGCGTTAACAGCCTTGCAATGCTCACCGCAAGTGGGATATTCAACTCATTGTACCCCGCAACCCTAAACTTCATGAAGGGCTTCCCTTTCTTGCTTGGGGCAGGTCTCCTTCTCATACCTGCCGTTCTCATAGGGATGCTTGAGAAGGCAGACCCCCATTTAGAGTTCCAACAATTCCCTCAATCACCCTGACTACTCTGGCGTCGATGAGGGA |

| | | |
|---|---|---|
| SLC10A6 | 1178 | CAATCCGCCCTCACTACAACCGATGAGAGCTAATTGTTC TTCTTCTTCTGCTTGTCCAGCTAATTCTTCTGAAGAAGA ACTGCCAGTGGGACTGGAAGTGCACGGAAATCTGGAAC TGGTGTTTACAGTGGTGTCTACAGTGATGATGGGACTGC TGATGTTTTCTCTGGGATGTTCTGTGGAAATTAGAAAAC TGTGGTCTCACATTAGAAGACCATGGGGAATTGCTGTGG GACTGCTGTGTCAGTTTGGACTGATGCCATTTACAGCTT ATCTGCTGGCTATTTCTTTTTCTCTGAAACCAGTGCAGG CTATTGCTGTGCTGATTATGGGATGTTGTCCAGGAGGAA CAATTTCTAATATTTTTACATTTTGGGTGGATGGAGATAT GGATCTGTCTATTTCTATGACAACATGTTCTACAGTGGCT GCTCTGGGAATGATGCCACTGTGTATTTATCTGTATACAT GGTCTTGGTCTCTGCAGCAGAATCTGACAATTCCATATC AGAATATTGGAATTACACTGGTGTGTCTGACAATTCCAG TGGCTTTTGGAGTGTATGTGAATTATAGATGGCCAAAAC AGTCTAAAATTATTCTGAAAATTGGAGCTGTGGTGGGAG GAGTGCTGCTGCTGGTGGTGGCTGTGGCTGGAGTGGTG CTGGCTAAAGGATCTTGGAATTCTGATATTACACTGCTG ACAATTTCTTTTATTTTTCCACTGATTGGACACGTGACAG GATTTCTGCTGGCTCTGTTTACACACCAGTCTTGGCAGA GATGTAGAACAATTTCTCTGGAAACAGGAGCTCAGAATA TTCAGATGTGTATTACAATGCTGCAGCTGTCTTTTACAGC TGAACACCTGGTGCAGATGCTGTCTTTTCCACTGGCTTA TGGACTGTTTCAGCTGATTGATGGATTTCTGATTGTGGCT GCTTATCAGACATATAAAAGAAGACTGAAAAATAAACAC GGAAAAAAAAATTCTGGATGTACAGAAGTGTGTCACAC AAGAAAATCTACATCTTCTAGAGAAACAAATGCTTTTCT GGAAGTGAATGAAGAAGGAGCTATTACACCAGGACCAC CAGGACCAATGGATTGTCACAGAGCTCTGGAACCAGTG GGACACATTACATCTTGTGAATAACTACTCTGGCGTCGA TGAGGGA |
| SLC16A10 | 1592 | CAATCCGCCCTCACTACAACCGATGGTGCTGTCTCAGGA AGAACCAGATTCTGCTAGAGGAACATCTGAAGCTCAGC CACTGGGACCAGCTCCAACAGGAGCTGCTCCACCACCA GGACCAGGACCATCTGATTCTCCAGAAGCTGCTGTGGA AAAAGTGGAAGTGGAACTGGCTGGACCAGCTACAGCTG AACCACACGAACCACCAGAACCACCAGAAGGAGGATG GGGATGGCTGGTGATGCTGGCTGCTATGTGGTGTAATGG ATCTGTGTTTGGAATTCAGAATGCTTGTGGAGTGCTGTT TGTGTCTATGCTGGAAACATTTGGATCTAAAGATGATGAT AAAATGGTGTTTAAAACAGCTTGGGTGGGATCTCTGTCT ATGGGAATGATTTTTTTTTGTTGTCCAATTGTGTCTGTGT TTACAGATCTGTTTGGATGTAGAAAAACAGCTGTGGTGG GAGCTGCTGTGGGATTTGTGGGACTGATGTCTTCTTCTT TTGTGTCTTCTATTGAACCACTGTATCTGACATATGGAAT TATTTTTGCTTGTGGATGTTCTTTTGCTTATCAGCCATCTC TGGTGATTCTGGGACACTATTTTAAAAAAAGACTGGGAC TGGTGAATGGAATTGTGACAGCTGGATCTTCTGTGTTTA CAATTCTGCTGCCACTGCTGCTGAGAGTGCTGATTGATT |

| | | |
|---|---|---|
| | | CTGTGGGACTGTTTTATACACTGAGAGTGCTGTGTATTTT<br>TATGTTTGTGCTGTTTCTGGCTGGATTTACATATAGACCA<br>CTGGCTACATCTACAAAAGATAAAGAATCTGGAGGATCT<br>GGATCTTCTCTGTTTTCTAGAAAAAAATTTTCTCCACCA<br>AAAAAAATTTTTAATTTTGCTATTTTTAAAGTGACAGCTT<br>ATGCTGTGTGGGCTGTGGGAATTCCACTGGCTCTGTTTG<br>GATATTTTGTGCCATATGTGCACCTGATGAAACACGTGA<br>ATGAAAGATTTCAGGATGAAAAAAATAAAGAAGTGGTG<br>CTGATGTGTATTGGAGTGACATCTGGAGTGGGAAGACTG<br>CTGTTTGGAAGAATTGCTGATTATGTGCCAGGAGTGAAA<br>AAAGTGTATCTGCAGGTGCTGTCTTTTTTTTTTATTGGAC<br>TGATGTCTATGATGATTCCACTGTGTTCTATTTTTGGAGC<br>TCTGATTGCTGTGTGTCTGATTATGGGACTGTTTGATGGA<br>TGTTTTATTTCTATTATGGCTCCAATTGCTTTTGAACTGGT<br>GGGAGCTCAGGATGTGTCTCAGGCTATTGGATTTCTGCT<br>GGGATTTATGTCTATTCCAATGACAGTGGGACCACCAAT<br>TGCTGGACTGCTGAGAGATAAACTGGGATCTTATGATGT<br>GGCTTTTTATCTGGCTGGAGTGCCACCACTGATTGGAGG<br>AGCTGTGCTGTGTTTTATTCCATGGATTCACTCTAAAAAA<br>CAGAGAGAAATTTCTAAAACAACAGGAAAAGAAAAAA<br>TGGAAAAAATGCTGGAAAATCAGAATTCTCTGCTGTCTT<br>CTTCTTCTGGAATGTTTAAAAAAGAATCTGATTCTATTAT<br>TTAACTACTCTGGCGTCGATGAGGGA |

**Supplementary Table 4.** List of potential transport events retained after LC-MS/MS data cleaning by manual peak inspection. Candidate transport-associated features were manually inspected for chromatographic peak quality, as illustrated in Figure S3. The retained potential transport events are listed by transporter, ionization mode (ESI+ or ESI-), and sample fraction in which they were detected (medium or oocyte extract). Metabolite annotations are putative unless independently confirmed by MS2 comparison with reference standards.

| CID | Name | Formula | Fold change | p-value | m/z | RT (min) | Annotation level | MS2 | Annotation note |
|---|---|---|---|---|---|---|---|---|---|
| **SLC10A2 (n = 73)** | | | | | | | | | |
| **ESI+ (n = 47)** | | | | | | | | | |
| **Medium (n = 9)** | | | | | | | | | |
| V4080 | (4R,5S,7R,11x)-11,12-Dihydroxy-1(10)-spirovetiven-2-one 12-glucoside | C21 H34 O8 | -1.204 | 5.307E-04 | 415.2331 | 7.600 | 3 | No MS2 | |
| V2913 | 1,2-Dihexanoyl-sn-glycero-3-phosphoethanolamine | C17 H34 N O8 P | 0.510 | 2.198E-04 | 412.2099 | 5.847 | 3 | DDA for preferred ion | |
| V882 | Cotinine | C10 H12 N2 O | 2.659 | 4.849E-02 | 177.1024 | 1.240 | 1 | DDA for preferred ion | |
| V570 | Glycerophospho-N-palmitoyl ethanolamine | C21 H44 N O7 P | -0.634 | 7.436E-03 | 454.2927 | 9.835 | 1 | DDA for preferred ion | |
| V1204 | N-(1-Deoxy-1-fructosyl)leucine | C12 H23 N O7 | -6.857 | 3.629E-02 | 294.1552 | 1.341 | 3 | DDA for preferred ion | |
| V621 | N-(1-Deoxy-1-fructosyl)leucine | C12 H23 N O7 | -1.586 | 3.714E-02 | 294.1552 | 1.456 | 3 | DDA for preferred ion | |
| V1689 | N-Acetylarylamine | C8 H9 N O | 2.490 | 4.491E-02 | 136.0757 | 1.273 | 3 | DDA for preferred ion | |
| V4308 | Sphingosine | C18 H37 N O2 | -0.535 | 1.406E-02 | 300.2902 | 9.089 | 3 | No MS2 | |
| V2675 | Urocanic acid | C6 H6 N2 O2 | -1.426 | 2.629E-02 | 139.0503 | 1.123 | 3 | DDA for preferred ion | |
| **Extract (n = 38)** | | | | | | | | | |
| V1355 | (±)-2-Methylthiazolidine | C4 H9 N S | 1.457 | 5.758E-04 | 104.0529 | 1.036 | 3 | DDA for preferred ion | |
| V2041 | 1-Benzylimidazole | C10 H10 N2 | 0.726 | 6.554E-03 | 159.0917 | 4.706 | 3 | DDA for preferred ion | |
| V1270 | 1-Naphthyl isocyanate | C11 H7 N O | 0.712 | 4.262E-03 | 170.0601 | 4.691 | 3 | DDA for preferred ion | |
| V3019 | 2,4,6-Octatriyn-1-ol | C8 H6 O | 1.417 | 3.712E-03 | 119.0491 | 1.223 | 3 | DDA for preferred ion | |
| V5067 | 2,5-Dioxopyrrolidin-1-yl 4-(bis(4-chlorophenyl)methyl)piperazine-1-carboxylate | C22 H21 Cl2 N3 O4 | 1.760 | 1.036E-02 | 462.0988 | 4.700 | 4 | No MS2 | |
| V4319 | 2-Octenoylcarnitine | C15 H27 N O4 | -1.619 | 3.254E-02 | 286.2017 | 6.748 | 3 | No MS2 | |
| V4530 | 3-[6-(2-methylpropyl)-2-oxo-1H-pyrazin-3-yl]propanamide | C11 H17 N3 O2 | 0.512 | 4.729E-02 | 224.1395 | 1.127 | 3 | DDA for preferred ion | |
| V4339 | 3-hydroxydecanoyl carnitine | C17 H33 N O5 | -0.634 | 4.127E-02 | 332.2436 | 6.980 | 3 | DDA for preferred ion | |
| V1313 | 3-Methylene-indolenine | C9 H7 N | 0.504 | 2.889E-03 | 130.0651 | 4.716 | 3 | DDA for preferred ion | |
| V4228 | 3-O-Methylniveusin A | C21 H28 O8 | 1.550 | 4.447E-02 | 409.1874 | 4.702 | 3 | No MS2 | |
| V637 | 6-Methylquinoline | C10 H9 N | 0.709 | 2.880E-03 | 144.0808 | 4.697 | 1 | DDA for preferred ion | |
| V1782 | 8-Methylundecanoylcarnitine | C19 H37 N O4 | -1.112 | 9.570E-03 | 344.2800 | 8.317 | 3 | DDA for preferred ion | |
| V3033 | Anabasine | C10 H14 N2 | 0.559 | 1.964E-02 | 163.1230 | 1.046 | 3 | No MS2 | |
| V5952 | Bezafibrate | C19 H20 Cl N O4 | 1.855 | 2.279E-02 | 362.1140 | 1.444 | 4 | No MS2 | |
| V2285 | Cinnamic acid | C9 H8 O2 | 1.262 | 1.065E-03 | 149.0597 | 2.652 | 3 | DDA for preferred ion | confirmed not cinnamic acid |
| V4150 | Deoxyadenosine | C10 H13 N5 O3 | 0.505 | 1.395E-02 | 252.1095 | 1.384 | 3 | No MS2 | |
| V5908 | Fluralaner | C22 H17 Cl2 F6 N3 O3 | 1.530 | 3.469E-05 | 556.0624 | 1.215 | 4 | No MS2 | |
| V3198 | gamma-Glutamylcysteine | C8 H14 N2 O5 S | 1.061 | 7.116E-03 | 251.0696 | 1.009 | 3 | DDA for other ion | |

| CID | Name | Formula | Fold change | p-value | m/z | RT (min) | Annotation level | MS2 | Annotation note |
|---|---|---|---|---|---|---|---|---|---|
| V4720 | gamma-Glutamylisoleucine | C11 H20 N2 O5 | 1.803 | 6.300E-04 | 261.1448 | 4.974 | 3 | DDA for preferred ion | |
| V5350 | Glutamyltryptophan | C16 H19 N3 O5 | 2.361 | 1.595E-03 | 334.1403 | 5.504 | 3 | No MS2 | |
| V3316 | Indene | C9 H8 | 0.728 | 3.939E-03 | 117.0698 | 4.702 | 3 | DDA for preferred ion | |
| V2443 | Indole | C8 H7 N | 0.739 | 6.144E-03 | 118.0652 | 1.243 | 3 | DDA for preferred ion | |
| V2619 | Indole | C8 H7 N | 0.982 | 6.188E-03 | 118.0652 | 2.665 | 3 | DDA for preferred ion | |
| V528 | Indole | C8 H7 N | 0.676 | 4.237E-03 | 118.0651 | 4.698 | 3 | DDA for preferred ion | |
| V50 | L-Phenylalanine | C9 H11 N O2 | 1.372 | 8.849E-04 | 166.0864 | 2.650 | 3 | DDA for preferred ion | confirmed by MS/MS spectral matching |
| V185 | L-Tryptophan | C11 H12 N2 O2 | 0.695 | 5.033E-03 | 205.0973 | 4.692 | 1 | DDA for preferred ion | confirmed by MS/MS spectral matching |
| V5106 | Mercaptopurine | C5 H4 N4 S | 0.556 | 4.936E-02 | 153.0236 | 5.332 | 3 | No MS2 | |
| V1172 | N-Acetylarylamine | C8 H9 N O | 1.357 | 3.281E-03 | 136.0757 | 1.214 | 3 | DDA for preferred ion | |
| V185 | N-Desmethyl Mephenytoin | C11 H12 N2 O2 | 0.695 | 5.033E-03 | 205.0973 | 4.692 | 1 | DDA for preferred ion | |
| V3980 | N-Docosahexaenoyl Threonine | C26 H39 N O4 | 2.195 | 7.854E-03 | 430.2956 | 7.514 | 3 | DDA for preferred ion | |
| V3733 | Ophthalmic acid | C11 H19 N3 O6 | 0.812 | 2.474E-02 | 290.1351 | 1.135 | 3 | No MS2 | |
| V4007 | Paracetamol sulfate | C8 H9 N O5 S | 2.019 | 1.254E-02 | 232.0269 | 1.488 | 3 | No MS2 | |
| V2451 | Trans-Non-2-en-(4.6.8)-triyn-1-ol | C9 H6 O | 1.437 | 6.890E-04 | 131.0492 | 2.640 | 3 | DDA for preferred ion | |
| V4654 | Tropolone A | C24 H33 N O6 | 0.591 | 2.569E-02 | 432.2386 | 8.862 | 3 | No MS2 | |
| V871 | Tyrosine fragement [M+H-NH3]+1 | C9 H8 O3 | 1.423 | 3.309E-03 | 165.0547 | 1.223 | 1 | DDA for preferred ion | confirmed by MS/MS spectral matching |
| V3180 | Uridine | C9 H12 N2 O6 | 2.831 | 2.081E-02 | 245.0771 | 1.224 | 3 | No MS2 | |
| V3209 | Uridine | C9 H12 N2 O6 | 1.781 | 1.411E-02 | 245.0772 | 1.172 | 3 | No MS2 | |
| V3212 | Uridine | C9 H12 N2 O6 | 1.781 | 1.411E-02 | 245.0771 | 1.197 | 3 | No MS2 | |
| **ESI- (n = 26)** | | | | | | | | | |
| **Medium (n = 2)** | | | | | | | | | |
| V554 | 1-(1,2,3,4,5-Pentahydroxypent-1-yl)-1,2,3,4-tetrahydro-beta-carboline-3-carboxylate | C17 H22 N2 O7 | -0.556 | 5.758E-03 | 365.1357 | 4.632 | 3 | No MS2 | |
| V49 | Sulfolithocholylglycine | C26 H43 N O7 S | -0.503 | 4.871E-02 | 512.2698 | 8.434 | 3 | DDA for preferred ion | |
| **Extract (n = 24)** | | | | | | | | | |
| V566 | (2S,3R)-3-Hydroxy-2-methylbutanoic acid | C5 H10 O3 | 1.730 | 1.325E-03 | 117.0553 | 4.064 | 3 | DDA for preferred ion | |
| V522 | 2-Hydroxy-3-methylpentanoic acid | C6 H12 O3 | 3.348 | 2.327E-05 | 131.0716 | 5.731 | 3 | DDA for preferred ion | |
| V512 | 2-Methyl-3-ketovaleric acid | C6 H10 O3 | 0.889 | 4.380E-04 | 129.0558 | 5.336 | 3 | DDA for preferred ion | |
| V513 | 2-Methyl-3-ketovaleric acid | C6 H10 O3 | 0.818 | 1.564E-03 | 129.0559 | 4.998 | 3 | DDA for preferred ion | |
| V474 | 3-Phenyllactic acid | C9 H10 O3 | 1.679 | 5.179E-05 | 165.0560 | 6.055 | 1 | DDA for preferred ion | confirmed by MS/MS spectral matching |
| V450 | 4-Hydroxystyrene | C8 H8 O | 2.039 | 4.724E-04 | 119.0503 | 6.052 | 3 | No MS2 | |
| V451 | 4-Hydroxystyrene | C8 H8 O | 2.085 | 1.103E-02 | 119.0501 | 4.721 | 3 | No MS2 | |
| V446 | 4-O-demethylhypothemycin | C18 H20 O8 | 3.892 | 6.334E-03 | 363.1072 | 1.322 | 4 | No MS2 | |
| V376 | AZT | C10 H13 N5 O4 | 4.785 | 3.424E-02 | 266.0896 | 1.469 | 3 | No MS2 | |
| V352 | Cinnamic acid | C9 H8 O2 | 1.905 | 8.661E-05 | 147.0453 | 6.054 | 3 | No MS2 | |
| V351 | cis-Aconitic acid | C6 H6 O6 | 1.105 | 1.832E-02 | 173.0092 | 1.308 | 3 | No MS2 | |

| CID | Name | Formula | Fold change | p-value | m/z | RT (min) | Annotation level | MS2 | Annotation note |
|---|---|---|---|---|---|---|---|---|---|
| V311 | Diphenyl sulfide | C12 H10 S | 2.604 | 2.095E-03 | 185.0434 | 4.065 | 3 | No MS2 | |
| V310 | DL-4-Hydroxyphenyllactic acid | C9 H10 O4 | 2.121 | 1.623E-02 | 181.0509 | 4.719 | 3 | DDA for preferred ion | |
| V307 | DOPA sulfate | C9 H11 N O7 S | 2.591 | 9.085E-05 | 276.0178 | 1.431 | 3 | DDA for preferred ion | |
| V276 | Geospallin B | C17 H24 N2 O7 S2 | 4.431 | 7.988E-04 | 431.0952 | 1.430 | 4 | No MS2 | |
| V273 | Glutamic acid glutamate | C10 H14 N2 O8 | 2.047 | 5.224E-03 | 289.0677 | 1.198 | 3 | DDA for preferred ion | |
| V266 | Glycocholic acid | C26 H43 N O6 | 2.889 | 3.728E-02 | 464.3027 | 7.576 | 1 | DDA for preferred ion | confirmed by MS/MS spectral matching |
| V238 | Kaempferol 3-p-coumarate | C24 H16 O8 | 2.870 | 3.340E-03 | 431.0763 | 2.626 | 4 | No MS2 | |
| V235 | L-Lactic acid | C3 H6 O3 | 1.119 | 1.132E-02 | 89.0247 | 1.026 | 4 | DDA for preferred ion | |
| V220 | L-Tyrosine | C9 H11 N O3 | 1.341 | 7.074E-03 | 180.0669 | 1.221 | 4 | DDA for preferred ion | |
| V177 | N-(3-Methyl-1,1-dioxo-1,4-thiazinan-4-yl)-1-(5-nitro-2-furanyl)methanimine | C10 H13 N3 O5 S | 0.674 | 3.276E-02 | 286.0505 | 1.377 | 3 | DDA for preferred ion | |
| V141 | Ophthalmic acid | C11 H19 N3 O6 | 0.818 | 1.933E-02 | 288.1203 | 1.155 | 3 | DDA for preferred ion | |
| V114 | Phenylacetic acid | C8 H8 O2 | 1.760 | 4.567E-03 | 135.0453 | 4.719 | 3 | No MS2 | |
| V21 | Uridine | C9 H12 N2 O6 | 1.990 | 3.856E-03 | 243.0622 | 1.199 | 1 | DDA for preferred ion | confirmed by MS/MS spectral matching |
| **SLC10A6 (n = 91)** | | | | | | | | | |
| **ESI+ (n = 37)** | | | | | | | | | |
| **Medium (n = 20)** | | | | | | | | | |
| V4080 | (4R,5S,7R,11x)-11,12-Dihydroxy-1(10)-spirovetiven-2-one 12-glucoside | C21 H34 O8 | -0.822 | 4.698E-02 | 415.2331 | 7.600 | 3 | No MS2 | |
| V1865 | (Z)-1,3-Octadiene | C8 H12 | -0.699 | 9.915E-04 | 109.1012 | 6.197 | 3 | DDA for preferred ion | |
| V2913 | 1,2-Dihexanoyl-sn-glycero-3-phosphoethanolamine | C17 H34 N O8 P | 0.576 | 1.100E-04 | 412.2099 | 5.847 | 3 | DDA for preferred ion | |
| V1340 | 1-(1,2,3,4,5-Pentahydroxypent-1-yl)-1,2,3,4-tetrahydro-beta-carboline-3-carboxylate | C17 H22 N2 O7 | -0.740 | 4.882E-03 | 367.1505 | 4.611 | 3 | DDA for preferred ion | |
| V5656 | 3',4',5'-Trimethoxycinnamyl alcohol acetate | C14 H18 O5 | -0.723 | 1.319E-04 | 267.1232 | 8.477 | 3 | No MS2 | |
| V1948 | Butenylcarnitine | C11 H19 N O4 | -1.578 | 3.304E-02 | 230.1390 | 1.339 | 3 | No MS2 | |
| V1013 | Butyrylcarnitine | C11 H21 N O4 | -0.525 | 3.030E-02 | 232.1546 | 3.853 | 3 | DDA for preferred ion | confirmed not butyrylcarnitine |
| V6026 | Chinikomycin B | C31 H35 Cl N2 O6 | -0.517 | 2.562E-02 | 567.2256 | 9.018 | 4 | No MS2 | |
| V4291 | Dihydroisomorphine-3-glucuronide | C23 H29 N O9 | -0.534 | 1.177E-02 | 464.1918 | 6.707 | 3 | No MS2 | |
| V4325 | Dodemorph | C18 H35 N O | -0.595 | 4.383E-02 | 282.2796 | 8.843 | 3 | No MS2 | |
| V4084 | Isoelemicin | C12 H16 O3 | -0.514 | 5.430E-03 | 209.1176 | 8.467 | 3 | No MS2 | |
| V5574 | Monoisobutyl phthalic acid | C12 H14 O4 | -0.545 | 2.074E-02 | 223.0968 | 7.627 | 3 | No MS2 | |
| V903 | N-(1-Deoxy-1-fructosyl)phenylalanine | C15 H21 N O7 | -0.782 | 9.796E-04 | 328.1397 | 2.810 | 3 | DDA for preferred ion | |
| V3824 | N-(2-Guanidinoethyl)-5-isoquinolinesulfonamide | C12 H15 N5 O2 S | -0.816 | 1.677E-02 | 294.1011 | 1.135 | 3 | No MS2 | |
| V4558 | Not named | C32 H39 N O7 | -2.156 | 2.704E-02 | 550.2805 | 9.791 | 3 | No MS2 | |
| V4478 | Phthalic Acid, Bis-Propyl Ester | C14 H18 O4 | -0.514 | 2.643E-02 | 251.1282 | 8.468 | 3 | No MS2 | |
| V4308 | Sphingosine | C18 H37 N O2 | -0.672 | 9.714E-03 | 300.2902 | 9.089 | 3 | No MS2 | |
| V661 | Sporovexin C | C15 H19 N O6 | -0.795 | 7.427E-04 | 310.1290 | 2.810 | 3 | DDA for preferred ion | |
| V5581 | Tetracosahexaenoic acid | C24 H36 O2 | -0.516 | 3.185E-02 | 357.2794 | 8.952 | 3 | No MS2 | |

| CID | Name | Formula | Fold change | p-value | m/z | RT (min) | Annotation level | MS2 | Annotation note |
|---|---|---|---|---|---|---|---|---|---|
| V3799 | Tetrahydrofuran | C4 H8 O | -0.892 | 5.015E-04 | 73.0648 | 8.008 | 3 | DDA for preferred ion | |
| **Extract (n = 17)** | | | | | | | | | |
| V3113 | (9E)-10-nitrooctadecenoic Acid | C18 H33 N O4 | -0.615 | 3.797E-02 | 328.2488 | 7.628 | 3 | DDA for preferred ion | |
| V1355 | (±)-2-Methylthiazolidine | C4 H9 N S | -0.953 | 4.206E-03 | 104.0529 | 1.036 | 3 | DDA for preferred ion | |
| V2041 | 1-Benzylimidazole | C10 H10 N2 | -1.138 | 2.010E-04 | 159.0917 | 4.706 | 3 | DDA for preferred ion | |
| V1270 | 1-Naphthyl isocyanate | C11 H7 N O | -1.202 | 8.894E-04 | 170.0601 | 4.691 | 3 | DDA for preferred ion | |
| V4339 | 3-hydroxydecanoyl carnitine | C17 H33 N O5 | -0.548 | 4.502E-02 | 332.2436 | 6.980 | 3 | DDA for preferred ion | |
| V3959 | 3-hydroxytridecanoyl carnitine | C20 H39 N O5 | -0.605 | 3.923E-03 | 374.2906 | 7.778 | 3 | No MS2 | |
| V1313 | 3-Methylene-indolenine | C9 H7 N | -0.765 | 1.536E-03 | 130.0651 | 4.716 | 3 | DDA for preferred ion | |
| V637 | 6-Methylquinoline | C10 H9 N | -1.104 | 4.691E-04 | 144.0808 | 4.697 | 1 | DDA for preferred ion | |
| V6004 | Cardinalin 8 | C32 H38 O10 | -1.489 | 1.296E-03 | 583.2552 | 7.656 | 3 | No MS2 | |
| V5908 | Fluralaner | C22 H17 Cl2 F6 N3 O3 | 0.555 | 6.751E-03 | 556.0624 | 1.215 | 4 | No MS2 | |
| V5350 | Glutamyltryptophan | C16 H19 N3 O5 | 0.987 | 2.146E-02 | 334.1403 | 5.504 | 3 | No MS2 | |
| V3316 | Indene | C9 H8 | -0.846 | 1.780E-04 | 117.0698 | 4.702 | 3 | DDA for preferred ion | |
| V528 | Indole | C8 H7 N | -1.085 | 7.041E-04 | 118.0651 | 4.698 | 3 | DDA for preferred ion | |
| V185 | L-Tryptophan | C11 H12 N2 O2 | -1.032 | 4.777E-04 | 205.0973 | 4.692 | 1 | DDA for preferred ion | confirmed by MS/MS spectral matching |
| V3828 | Naphthalene epoxide | C10 H8 O | -0.904 | 3.760E-04 | 145.0648 | 4.699 | 4 | DDA for preferred ion | |
| V2373 | O-Propanoyl-D-carnitine | C10 H19 N O4 | -1.764 | 2.627E-04 | 218.1390 | 1.671 | 3 | DDA for preferred ion | |
| V585 | Tridec-3-enoylcarnitine | C20 H37 N O4 | -0.522 | 7.123E-03 | 356.2798 | 8.303 | 3 | DDA for preferred ion | |
| **ESI- (n = 54)** | | | | | | | | | |
| **Medium (n = 44)** | | | | | | | | | |
| V572 | (-)-Nopol | C11 H18 O | -1.076 | 6.518E-03 | 165.1288 | 7.486 | 3 | No MS2 | |
| V566 | (2S,3R)-3-Hydroxy-2-methylbutanoic acid | C5 H10 O3 | -0.691 | 1.371E-02 | 117.0553 | 4.064 | 3 | DDA for preferred ion | |
| V565 | (2xi,6xi)-7-Methyl-3-methylene-1,2,6,7-octanetetrol | C10 H20 O4 | -0.583 | 3.357E-02 | 203.1291 | 6.562 | 3 | No MS2 | |
| V557 | (Z)-3-Methyl-3-decenoic acid | C11 H20 O2 | -0.882 | 1.476E-02 | 183.1393 | 7.484 | 3 | DDA for preferred ion | |
| V554 | 1-(1,2,3,4,5-Pentahydroxypent-1-yl)-1,2,3,4-tetrahydro-beta-carboline-3-carboxylate | C17 H22 N2 O7 | -0.683 | 2.286E-03 | 365.1357 | 4.632 | 3 | No MS2 | |
| V549 | 1-Octen-3-yl glucoside | C14 H26 O6 | -0.546 | 1.778E-02 | 289.1661 | 6.995 | 3 | No MS2 | |
| V548 | 10,11-epoxychlorovulone | C21 H29 Cl O5 | -0.610 | 3.458E-02 | 395.1634 | 7.006 | 3 | No MS2 | |
| V544 | 13-HODE | C18 H32 O3 | -0.790 | 1.606E-02 | 295.2281 | 9.650 | 3 | DDA for preferred ion | |
| V537 | 2,2,4,4-Tetramethyl-6-(1-oxobutyl)-1,3,5-cyclohexanetrione | C14 H20 O4 | -1.068 | 7.188E-03 | 251.1292 | 7.717 | 3 | DDA for preferred ion | |
| V529 | 2-(7'-methylthio)heptylmalate | C12 H20 O5 S | 1.431 | 3.723E-02 | 275.0962 | 6.546 | 3 | DDA for preferred ion | |
| V507 | 2-Pentanamido-3-phenylpropanoic acid | C14 H19 N O3 | -0.505 | 1.020E-02 | 248.1295 | 7.332 | 3 | No MS2 | |
| V504 | 3'-Hydroxyamobarbital | C11 H18 N2 O4 | -0.640 | 9.186E-03 | 241.1195 | 5.677 | 3 | No MS2 | |
| V488 | 3-Hydroxycapric acid | C10 H20 O3 | -0.533 | 3.463E-03 | 187.1343 | 8.341 | 3 | DDA for preferred ion | |
| V481 | 3-Methylindole | C9 H9 N | -0.780 | 4.030E-02 | 130.0664 | 2.881 | 3 | No MS2 | |

| CID | Name | Formula | Fold change | p-value | m/z | RT (min) | Annotation level | MS2 | Annotation note |
|---|---|---|---|---|---|---|---|---|---|
| V467 | 4,11,13,15-Tetrahydroridentin B | C15 H24 O4 | -1.408 | 4.614E-03 | 267.1605 | 8.290 | 3 | DDA for preferred ion | |
| V431 | 5-Hydroxy-N-formylkynurenine | C11 H12 N2 O5 | -1.024 | 3.664E-02 | 251.0675 | 4.860 | 3 | No MS2 | |
| V420 | 7-Hydroxyoctanoic acid | C8 H16 O3 | -0.545 | 7.811E-03 | 159.1030 | 7.282 | 3 | No MS2 | |
| V419 | 8-Hydroxymianserin | C18 H20 N2 O | -0.694 | 4.419E-02 | 279.1506 | 9.430 | 4 | DDA for preferred ion | |
| V407 | Acetyl-N-formyl-5-methoxykynurenamine | C13 H16 N2 O4 | -0.525 | 1.051E-02 | 263.1039 | 5.442 | 3 | DDA for preferred ion | |
| V375 | Benzocaine | C9 H11 N O2 | -0.695 | 1.490E-03 | 164.0719 | 2.855 | 3 | No MS2 | |
| V369 | Bisphenol S | C12 H10 O4 S | -0.604 | 3.966E-02 | 249.0227 | 6.568 | 3 | No MS2 | |
| V363 | Cabbage identification factor 2 | C15 H12 N4 O3 S | -0.800 | 6.913E-03 | 327.0542 | 7.485 | 4 | No MS2 | |
| V362 | Carbofuran, 3OH- | C12 H15 N O4 | -0.710 | 1.820E-06 | 236.0930 | 2.850 | 3 | No MS2 | |
| V338 | Cortolone-3-glucuronide | C27 H42 O11 | -0.569 | 8.607E-03 | 541.2668 | 6.562 | 3 | No MS2 | |
| V302 | Enterolactone 3'-sulfate | C18 H18 O7 S | -1.629 | 3.883E-02 | 377.0697 | 9.536 | 4 | No MS2 | |
| V301 | Eremopetasinorol | C13 H20 O2 | -1.081 | 6.342E-03 | 207.1393 | 7.717 | 3 | DDA for preferred ion | |
| V283 | Gallicynoic acid F | C18 H32 O6 | -1.016 | 1.254E-02 | 343.2129 | 6.770 | 3 | No MS2 | |
| V276 | Geospallin B | C17 H24 N2 O7 S2 | -0.659 | 2.741E-02 | 431.0952 | 1.430 | 4 | No MS2 | |
| V265 | Glycoursodeoxycholic acid | C26 H43 N O5 | -0.543 | 1.246E-03 | 448.3076 | 8.367 | 1 | DDA for preferred ion | |
| V249 | Indole-3-carboxaldehyde | C9 H7 N O | -0.558 | 3.149E-02 | 144.0457 | 6.419 | 3 | DDA for preferred ion | |
| V238 | Kaempferol 3-p-coumarate | C24 H16 O8 | -0.733 | 2.705E-02 | 431.0763 | 2.626 | 4 | No MS2 | |
| V225 | L-Isoleucine | C6 H13 N O2 | -0.927 | 7.683E-03 | 130.0876 | 1.476 | 3 | DDA for preferred ion | |
| V216 | Leukotriene B4 | C20 H32 O4 | -0.545 | 9.532E-03 | 335.2231 | 8.834 | 3 | DDA for preferred ion | |
| V196 | Methylone | C11 H13 N O3 | -0.557 | 2.493E-04 | 206.0825 | 6.096 | 3 | DDA for preferred ion | |
| V181 | N-(1-Deoxy-1-fructosyl)leucine | C12 H23 N O7 | -0.707 | 3.329E-03 | 292.1405 | 1.370 | 3 | DDA for preferred ion | |
| V167 | N-Desmethyl Mephenytoin | C11 H12 N2 O2 | -0.550 | 1.317E-02 | 203.0826 | 4.928 | 3 | No MS2 | |
| V165 | N-Lactoylleucine | C9 H17 N O4 | -1.048 | 4.020E-03 | 202.1088 | 1.495 | 3 | DDA for preferred ion | |
| V96 | Prehumulinic acid | C16 H24 O4 | -1.332 | 4.433E-03 | 279.1605 | 8.330 | 3 | No MS2 | |
| V94 | Prolylhydroxyproline | C10 H16 N2 O4 | -1.089 | 1.070E-02 | 227.1037 | 4.813 | 3 | No MS2 | |
| V49 | Sulfolithocholylglycine | C26 H43 N O7 S | -0.593 | 2.175E-03 | 512.2698 | 8.434 | 3 | DDA for preferred ion | |
| V43 | Tetrahydrofurfuryl butyrate | C9 H16 O3 | -0.709 | 4.317E-02 | 171.1030 | 5.764 | 3 | DDA for preferred ion | |
| V44 | Tetrahydrofurfuryl butyrate | C9 H16 O3 | -0.535 | 1.075E-02 | 171.1029 | 6.835 | 3 | DDA for preferred ion | |
| V37 | Traumatic acid | C12 H20 O4 | -0.853 | 8.354E-03 | 227.1292 | 7.485 | 3 | DDA for preferred ion | |
| V27 | Ubiquinone-2 | C19 H26 O4 | -0.907 | 1.291E-02 | 317.1761 | 8.760 | 3 | No MS2 | |
| **Extract (n = 10)** | | | | | | | | | |
| V566 | (2S,3R)-3-Hydroxy-2-methylbutanoic acid | C5 H10 O3 | 0.967 | 1.837E-02 | 117.0553 | 4.064 | 3 | DDA for preferred ion | |
| V458 | 4-Hydroxy-2-butenoic acid gamma-lactone | C4 H4 O2 | 0.762 | 4.669E-02 | 83.0140 | 1.873 | 3 | No MS2 | |
| V433 | 5-Hydroxy-2-furoic acid | C5 H4 O4 | 0.635 | 3.510E-02 | 127.0038 | 1.865 | 3 | No MS2 | |
| V357 | Chloramphenicol | C11 H12 Cl2 N2 O5 | 2.018 | 2.378E-02 | 321.0052 | 6.706 | 3 | No MS2 | |

| CID | Name | Formula | Fold change | p-value | m/z | RT (min) | Annotation level | MS2 | Annotation note |
|---|---|---|---|---|---|---|---|---|---|
| V325 | D-4'-Phosphopantothenate | C9 H18 N O8 P | -0.501 | 1.994E-02 | 298.0698 | 1.514 | 3 | No MS2 | |
| V278 | gamma-Glutamylisoleucine | C11 H20 N2 O5 | 1.243 | 5.528E-03 | 259.1301 | 5.419 | 3 | No MS2 | |
| V235 | L-Lactic acid | C3 H6 O3 | 1.479 | 3.173E-03 | 89.0247 | 1.026 | 4 | DDA for preferred ion | |
| V182 | N'-Formylkynurenine | C11 H12 N2 O4 | -1.334 | 3.818E-02 | 235.0726 | 2.873 | 3 | DDA for preferred ion | |
| V136 | Oxoadipic acid | C6 H8 O5 | 0.887 | 4.955E-02 | 159.0301 | 1.877 | 3 | DDA for preferred ion | |
| V42 | Thymidine | C10 H14 N2 O5 | -0.904 | 3.784E-03 | 241.0832 | 2.856 | 3 | DDA for preferred ion | |
| **SLC13A2 (n = 33)** | | | | | | | | | |
| **ESI+ (n = 15)** | | | | | | | | | |
| **Medium (n = 5)** | | | | | | | | | |
| V1797 | Guanine | C5 H5 N5 O | 4.868 | 4.541E-02 | 152.0567 | 1.330 | 3 | DDA for preferred ion | |
| V259 | Hexanoylcarnitine | C13 H25 N O4 | 1.585 | 3.219E-02 | 260.1857 | 5.831 | 1 | DDA for preferred ion | confirmed by MS/MS spectral matching |
| V4238 | Methandienone | C20 H28 O2 | -0.605 | 1.357E-02 | 301.2168 | 8.804 | 3 | No MS2 | |
| V2929 | Morphine | C17 H19 N O3 | -0.604 | 1.966E-02 | 286.1443 | 8.599 | 3 | DDA for preferred ion | |
| V297 | Pantothenate | C9 H17 N O5 | 1.420 | 4.982E-02 | 220.1182 | 3.638 | 1 | DDA for preferred ion | |
| **Extract (n = 10)** | | | | | | | | | |
| V3019 | 2,4,6-Octatriyn-1-ol | C8 H6 O | 0.783 | 2.147E-02 | 119.0491 | 1.223 | 3 | DDA for preferred ion | |
| V5899 | 3-{4-[(6,7-Dimethoxyquinolin-4-yl)oxy]-3-fluorophenyl}-1-(2-phenylacetyl)thiourea | C26 H22 F N3 O4 S | -0.721 | 1.704E-02 | 492.1406 | 3.702 | 4 | No MS2 | |
| V4329 | Androsta-4,16-dien-3-one | C19 H26 O | -0.991 | 6.034E-03 | 271.2060 | 7.569 | 3 | No MS2 | |
| V1013 | Butyrylcarnitine | C11 H21 N O4 | 2.609 | 4.123E-02 | 232.1546 | 3.853 | 3 | DDA for preferred ion | confirmed not butyrylcarnitine |
| V5908 | Fluralaner | C22 H17 Cl2 F6 N3 O3 | 1.137 | 1.955E-04 | 556.0624 | 1.215 | 4 | No MS2 | |
| V4841 | Fradcarbazole C | C29 H25 N5 O3 | -1.030 | 4.565E-03 | 492.2011 | 5.538 | 4 | No MS2 | |
| V3198 | gamma-Glutamylcysteine | C8 H14 N2 O5 S | 1.627 | 3.061E-04 | 251.0696 | 1.009 | 3 | DDA for other ion | |
| V5350 | Glutamyltryptophan | C16 H19 N3 O5 | 1.510 | 4.083E-03 | 334.1403 | 5.504 | 3 | No MS2 | |
| V1172 | N-Acetylarylamine | C8 H9 N O | 0.756 | 2.148E-02 | 136.0757 | 1.214 | 3 | DDA for preferred ion | |
| V871 | Tyrosine fragement [M+H-NH3]+1 | C9 H8 O3 | 0.794 | 2.373E-02 | 165.0547 | 1.223 | 1 | DDA for preferred ion | confirmed by MS/MS spectral matching |
| **ESI- (n = 18)** | | | | | | | | | |
| **Medium (n = 3)** | | | | | | | | | |
| V554 | 1-(1,2,3,4,5-Pentahydroxypent-1-yl)-1,2,3,4-tetrahydro-beta-carboline-3-carboxylate | C17 H22 N2 O7 | -0.806 | 4.971E-02 | 365.1357 | 4.632 | 3 | No MS2 | |
| V255 | Hypoxanthine | C5 H4 N4 O | -4.393 | 4.488E-02 | 135.0312 | 1.114 | 1 | DDA for preferred ion | confirmed by MS/MS spectral matching |
| V42 | Thymidine | C10 H14 N2 O5 | 2.238 | 3.017E-02 | 241.0832 | 2.856 | 3 | DDA for preferred ion | |
| **Extract (n = 15)** | | | | | | | | | |
| V526 | 2-Aminobicyclo[3.1.0]hexane-2,6-dicarboxylic acid | C8 H11 N O4 | 1.223 | 4.056E-02 | 184.0617 | 4.607 | 3 | No MS2 | |
| V522 | 2-Hydroxy-3-methylpentanoic acid | C6 H12 O3 | 0.538 | 2.694E-02 | 131.0716 | 5.731 | 3 | DDA for preferred ion | |
| V474 | 3-Phenyllactic acid | C9 H10 O3 | -0.859 | 1.011E-02 | 165.0560 | 6.055 | 1 | DDA for preferred ion | confirmed by MS/MS spectral matching |
| V458 | 4-Hydroxy-2-butenoic acid gamma-lactone | C4 H4 O2 | -0.929 | 2.495E-02 | 83.0140 | 1.873 | 3 | No MS2 | |

| CID | Name | Formula | Fold change | p-value | m/z | RT (min) | Annotation level | MS2 | Annotation note |
|---|---|---|---|---|---|---|---|---|---|
| V433 | 5-Hydroxy-2-furoic acid | C5 H4 O4 | -0.909 | 9.041E-03 | 127.0038 | 1.865 | 3 | No MS2 | |
| V420 | 7-Hydroxyoctanoic acid | C8 H16 O3 | 0.804 | 2.818E-02 | 159.1030 | 7.282 | 3 | No MS2 | |
| V399 | Alpha-dihydroartemisinin | C15 H24 O5 | 1.009 | 9.738E-03 | 283.1553 | 7.415 | 3 | No MS2 | |
| V352 | Cinnamic acid | C9 H8 O2 | -0.654 | 1.602E-02 | 147.0453 | 6.054 | 3 | No MS2 | |
| V308 | DL-Tryptophan | C11 H12 N2 O2 | -4.061 | 2.813E-02 | 203.0828 | 4.689 | 1 | DDA for preferred ion | confirmed by MS/MS spectral matching |
| V280 | gamma-Glutamyl-S-(1-propenyl)cysteine sulfoxide | C11 H18 N2 O6 S | -3.955 | 1.045E-02 | 305.0814 | 4.518 | 3 | No MS2 | |
| V252 | Indole | C8 H7 N | -1.680 | 3.621E-02 | 116.0509 | 4.689 | 3 | DDA for preferred ion | |
| V220 | L-Tyrosine | C9 H11 N O3 | 0.745 | 2.910E-02 | 180.0669 | 1.221 | 4 | DDA for preferred ion | |
| V141 | Ophthalmic acid | C11 H19 N3 O6 | 0.771 | 3.228E-02 | 288.1203 | 1.155 | 3 | DDA for preferred ion | |
| V136 | Oxoadipic acid | C6 H8 O5 | -1.272 | 1.666E-02 | 159.0301 | 1.877 | 3 | DDA for preferred ion | |
| V45 | Tetrahydrodipicolinate | C7 H9 N O4 | 1.681 | 3.588E-02 | 170.0461 | 1.581 | 3 | No MS2 | |
| **SLC16A10 (n = 59)** | | | | | | | | | |
| **ESI+ (n = 27)** | | | | | | | | | |
| **Medium (n = 11)** | | | | | | | | | |
| V2236 | 6-Methyltetrahydropterin | C7 H11 N5 O | 0.983 | 1.207E-02 | 182.1037 | 1.009 | 3 | No MS2 | |
| V5070 | Biopterin-a-glucoside | C15 H21 N5 O8 | 2.132 | 1.442E-02 | 400.1461 | 8.307 | 4 | No MS2 | |
| V4393 | Dasabuvir | C26 H27 N3 O5 S | 1.942 | 1.243E-02 | 494.1728 | 8.016 | 4 | No MS2 | |
| V4666 | Ethyl vanillin isobutyrate | C13 H16 O4 | 0.581 | 2.386E-02 | 237.1125 | 9.073 | 3 | No MS2 | |
| V15 | Hypoxanthine | C5 H4 N4 O | 0.944 | 6.671E-04 | 137.0458 | 1.004 | 1 | DDA for preferred ion | |
| V700 | Imidazooxazole | C4 H3 N3 O | 0.961 | 1.410E-03 | 110.0349 | 1.002 | 3 | DDA for preferred ion | |
| V4238 | Methandienone | C20 H28 O2 | -0.601 | 1.734E-02 | 301.2168 | 8.804 | 3 | No MS2 | |
| V903 | N-(1-Deoxy-1-fructosyl)phenylalanine | C15 H21 N O7 | -0.611 | 2.491E-02 | 328.1397 | 2.810 | 3 | DDA for preferred ion | |
| V4318 | SCD1 Inhibitor | C21 H20 F3 N3 O3 | 1.954 | 3.088E-02 | 420.1538 | 7.379 | 4 | No MS2 | |
| V661 | Sporovexin C | C15 H19 N O6 | -0.601 | 2.933E-02 | 310.1290 | 2.810 | 3 | DDA for preferred ion | |
| V4877 | Stachybotrin A | C23 H31 N O5 | 0.523 | 2.228E-02 | 402.2280 | 9.072 | 3 | No MS2 | |
| **Extract (n = 16)** | | | | | | | | | |
| V1355 | (±)-2-Methylthiazolidine | C4 H9 N S | -1.462 | 8.903E-03 | 104.0529 | 1.036 | 3 | DDA for preferred ion | |
| V2041 | 1-Benzylimidazole | C10 H10 N2 | -2.706 | 2.514E-03 | 159.0917 | 4.706 | 3 | DDA for preferred ion | |
| V1702 | 1-Methylguanine | C6 H7 N5 O | -0.590 | 4.766E-04 | 166.0724 | 1.169 | 2 | DDA for preferred ion | |
| V1270 | 1-Naphthyl isocyanate | C11 H7 N O | -3.091 | 4.439E-03 | 170.0601 | 4.691 | 3 | DDA for preferred ion | |
| V3019 | 2,4,6-Octatriyn-1-ol | C8 H6 O | -1.482 | 9.084E-03 | 119.0491 | 1.223 | 3 | DDA for preferred ion | |
| V1313 | 3-Methylene-indolenine | C9 H7 N | -2.237 | 4.142E-03 | 130.0651 | 4.716 | 3 | DDA for preferred ion | |
| V637 | 6-Methylquinoline | C10 H9 N | -2.658 | 3.010E-03 | 144.0808 | 4.697 | 1 | DDA for preferred ion | |
| V2285 | Cinnamic acid | C9 H8 O2 | -0.758 | 5.600E-03 | 149.0597 | 2.652 | 3 | DDA for preferred ion | confirmed not cinnamic acid |
| V3316 | Indene | C9 H8 | -1.612 | 8.173E-04 | 117.0698 | 4.702 | 3 | DDA for preferred ion | |

| CID | Name | Formula | Fold change | p-value | m/z | RT (min) | Annotation level | MS2 | Annotation note |
|---|---|---|---|---|---|---|---|---|---|
| V528 | Indole | C8 H7 N | -2.517 | 3.400E-03 | 118.0651 | 4.698 | 3 | DDA for preferred ion | |
| V50 | L-Phenylalanine | C9 H11 N O2 | -1.118 | 3.424E-03 | 166.0864 | 2.650 | 3 | DDA for preferred ion | confirmed by MS/MS spectral matching |
| V185 | L-Tryptophan | C11 H12 N2 O2 | -2.454 | 2.772E-03 | 205.0973 | 4.692 | 1 | DDA for preferred ion | confirmed by MS/MS spectral matching |
| V3828 | Naphthalene epoxide | C10 H8 O | -1.531 | 1.772E-04 | 145.0648 | 4.699 | 4 | DDA for preferred ion | |
| V2373 | O-Propanoyl-D-carnitine | C10 H19 N O4 | -1.419 | 1.144E-03 | 218.1390 | 1.671 | 3 | DDA for preferred ion | |
| V2451 | Trans-Non-2-en-(4.6.8)-triyn-1-ol | C9 H6 O | -1.180 | 1.895E-03 | 131.0492 | 2.640 | 3 | DDA for preferred ion | |
| V871 | Tyrosine fragement [M+H-NH3]+1 | C9 H8 O3 | -1.479 | 8.074E-03 | 165.0547 | 1.223 | 1 | DDA for preferred ion | confirmed by MS/MS spectral matching |
| **ESI- (n = 32)** | | | | | | | | | |
| **Medium (n = 17)** | | | | | | | | | |
| V572 | (-)-Nopol | C11 H18 O | -0.527 | 5.716E-03 | 165.1288 | 7.486 | 3 | No MS2 | |
| V558 | (S)-9-Hydroxy-10-undecenoic acid | C11 H20 O3 | -0.937 | 3.849E-02 | 199.1344 | 6.538 | 3 | No MS2 | |
| V554 | 1-(1,2,3,4,5-Pentahydroxypent-1-yl)-1,2,3,4-tetrahydro-beta-carboline-3-carboxylate | C17 H22 N2 O7 | -0.870 | 2.936E-03 | 365.1357 | 4.632 | 3 | No MS2 | |
| V537 | 2,2,4,4-Tetramethyl-6-(1-oxobutyl)-1,3,5-cyclohexanetrione | C14 H20 O4 | -0.560 | 7.442E-03 | 251.1292 | 7.717 | 3 | DDA for preferred ion | |
| V507 | 2-Pentanamido-3-phenylpropanoic acid | C14 H19 N O3 | -0.884 | 2.188E-02 | 248.1295 | 7.332 | 3 | No MS2 | |
| V469 | 4'-Methoxychalcone | C16 H14 O2 | 2.391 | 3.402E-02 | 237.0911 | 1.194 | 3 | No MS2 | |
| V467 | 4,11,13,15-Tetrahydroridentin B | C15 H24 O4 | -0.597 | 4.896E-03 | 267.1605 | 8.290 | 3 | DDA for preferred ion | |
| V399 | Alpha-dihydroartemisinin | C15 H24 O5 | -1.135 | 7.102E-03 | 283.1553 | 7.415 | 3 | No MS2 | |
| V384 | Anhydroecgonine Methyl Ester | C10 H15 N O2 | -0.730 | 2.536E-02 | 180.1033 | 8.039 | 3 | No MS2 | |
| V301 | Eremopetasinorol | C13 H20 O2 | -0.565 | 4.776E-02 | 207.1393 | 7.717 | 3 | DDA for preferred ion | |
| V283 | Gallicynoic acid F | C18 H32 O6 | -0.733 | 1.658E-02 | 343.2129 | 6.770 | 3 | No MS2 | |
| V254 | Hypoxanthine | C5 H4 N4 O | 1.219 | 1.518E-04 | 135.0313 | 1.015 | 1 | DDA for preferred ion | confirmed by MS/MS spectral matching |
| V203 | Mebutamate | C10 H20 N2 O4 | -1.193 | 2.102E-02 | 231.1352 | 5.764 | 3 | DDA for preferred ion | |
| V196 | Methylone | C11 H13 N O3 | -0.668 | 9.892E-03 | 206.0825 | 6.096 | 3 | DDA for preferred ion | |
| V96 | Prehumulinic acid | C16 H24 O4 | -0.503 | 2.515E-02 | 279.1605 | 8.330 | 3 | No MS2 | |
| V43 | Tetrahydrofurfuryl butyrate | C9 H16 O3 | -1.138 | 1.710E-02 | 171.1030 | 5.764 | 3 | DDA for preferred ion | |
| V7 | Xylaric acid A | C14 H22 O5 | -1.777 | 1.189E-02 | 269.1396 | 7.203 | 3 | No MS2 | |
| **Extract (n = 15)** | | | | | | | | | |
| V484 | 3-Indoleacetic Acid | C10 H9 N O2 | -1.382 | 1.703E-02 | 174.0563 | 2.879 | 4 | No MS2 | |
| V481 | 3-Methylindole | C9 H9 N | -4.078 | 4.509E-02 | 130.0664 | 2.881 | 3 | No MS2 | |
| V474 | 3-Phenyllactic acid | C9 H10 O3 | -2.027 | 1.039E-02 | 165.0560 | 6.055 | 1 | DDA for preferred ion | confirmed by MS/MS spectral matching |
| V450 | 4-Hydroxystyrene | C8 H8 O | -1.580 | 4.153E-02 | 119.0503 | 6.052 | 3 | No MS2 | |
| V451 | 4-Hydroxystyrene | C8 H8 O | -1.871 | 4.949E-02 | 119.0501 | 4.721 | 3 | No MS2 | |
| V352 | Cinnamic acid | C9 H8 O2 | -1.711 | 4.731E-03 | 147.0453 | 6.054 | 3 | No MS2 | |
| V310 | DL-4-Hydroxyphenyllactic acid | C9 H10 O4 | -2.431 | 1.022E-02 | 181.0509 | 4.719 | 3 | DDA for preferred ion | |
| V308 | DL-Tryptophan | C11 H12 N2 O2 | -3.695 | 4.174E-02 | 203.0828 | 4.689 | 1 | DDA for preferred ion | confirmed by MS/MS spectral matching |
| V278 | gamma-Glutamylisoleucine | C11 H20 N2 O5 | 0.878 | 1.956E-02 | 259.1301 | 5.419 | 3 | No MS2 | |

| CID | Name | Formula | Fold change | p-value | m/z | RT (min) | Annotation level | MS2 | Annotation note |
|---|---|---|---|---|---|---|---|---|---|
| V260 | Hippuric acid | C9 H9 N O3 | -1.418 | 1.736E-02 | 178.0512 | 5.378 | 1 | DDA for preferred ion | confirmed by MS/MS spectral matching |
| V259 | Homogentisic acid | C8 H8 O4 | -0.881 | 1.339E-02 | 167.0351 | 1.501 | 3 | No MS2 | |
| V249 | Indole-3-carboxaldehyde | C9 H7 N O | -0.704 | 4.073E-02 | 144.0457 | 6.419 | 3 | DDA for preferred ion | |
| V250 | Indole-3-carboxaldehyde | C9 H7 N O | -1.098 | 8.250E-03 | 144.0457 | 2.873 | 3 | No MS2 | |
| V200 | Methyl 2,6-dihydroxy-4-quinolinecarboxylate | C11 H9 N O4 | -2.279 | 4.926E-02 | 218.0460 | 2.867 | 3 | No MS2 | |
| V182 | N'-Formylkynurenine | C11 H12 N2 O4 | -2.378 | 1.252E-02 | 235.0726 | 2.873 | 3 | DDA for preferred ion | |
| **SLC46A1 (n = 36)** | | | | | | | | | |
| **ESI+ (n = 13)** | | | | | | | | | |
| **Medium (n = 7)** | | | | | | | | | |
| V1865 | (Z)-1,3-Octadiene | C8 H12 | -0.506 | 3.125E-02 | 109.1012 | 6.197 | 3 | DDA for preferred ion | |
| V882 | Cotinine | C10 H12 N2 O | 2.539 | 4.694E-02 | 177.1024 | 1.240 | 1 | DDA for preferred ion | |
| V259 | Hexanoylcarnitine | C13 H25 N O4 | 1.989 | 9.295E-03 | 260.1857 | 5.831 | 1 | DDA for preferred ion | confirmed by MS/MS spectral matching |
| V297 | Pantothenate | C9 H17 N O5 | 1.794 | 1.848E-02 | 220.1182 | 3.638 | 1 | DDA for preferred ion | |
| V3180 | Uridine | C9 H12 N2 O6 | 1.454 | 3.501E-02 | 245.0771 | 1.224 | 3 | No MS2 | |
| V3212 | Uridine | C9 H12 N2 O6 | 1.416 | 3.533E-02 | 245.0771 | 1.197 | 3 | No MS2 | |
| V2675 | Urocanic acid | C6 H6 N2 O2 | -1.641 | 1.625E-02 | 139.0503 | 1.123 | 3 | DDA for preferred ion | |
| **Extract (n = 6)** | | | | | | | | | |
| V4319 | 2-Octenoylcarnitine | C15 H27 N O4 | -1.037 | 1.214E-02 | 286.2017 | 6.748 | 3 | No MS2 | |
| V5895 | 5-Methyldodecanoylcarnitine | C20 H39 N O4 | 0.518 | 9.705E-03 | 358.2959 | 8.475 | 3 | No MS2 | |
| V6004 | Cardinalin 8 | C32 H38 O10 | -1.077 | 1.050E-03 | 583.2552 | 7.656 | 3 | No MS2 | |
| V5908 | Fluralaner | C22 H17 Cl2 F6 N3 O3 | 0.684 | 5.670E-03 | 556.0624 | 1.215 | 4 | No MS2 | |
| V4841 | Fradcarbazole C | C29 H25 N5 O3 | -1.047 | 4.285E-02 | 492.2011 | 5.538 | 4 | No MS2 | |
| V5350 | Glutamyltryptophan | C16 H19 N3 O5 | 2.655 | 5.285E-04 | 334.1403 | 5.504 | 3 | No MS2 | |
| **ESI- (n = 23)** | | | | | | | | | |
| **Medium (n = 20)** | | | | | | | | | |
| V572 | (-)-Nopol | C11 H18 O | -1.097 | 6.529E-03 | 165.1288 | 7.486 | 3 | No MS2 | |
| V557 | (Z)-3-Methyl-3-decenoic acid | C11 H20 O2 | -0.865 | 1.221E-02 | 183.1393 | 7.484 | 3 | DDA for preferred ion | |
| V554 | 1-(1,2,3,4,5-Pentahydroxypent-1-yl)-1,2,3,4-tetrahydro-beta-carboline-3-carboxylate | C17 H22 N2 O7 | -0.735 | 4.500E-02 | 365.1357 | 4.632 | 3 | No MS2 | |
| V537 | 2,2,4,4-Tetramethyl-6-(1-oxobutyl)-1,3,5-cyclohexanetrione | C14 H20 O4 | -1.027 | 5.451E-03 | 251.1292 | 7.717 | 3 | DDA for preferred ion | |
| V533 | 2-(1,3-Dioxo-1,3-dihydro-isoindol-2-yl)-3-(1H-indol-3-yl)-propionic acid | C19 H14 N2 O4 | -0.535 | 2.412E-02 | 333.0884 | 9.393 | 3 | No MS2 | |
| V467 | 4,11,13,15-Tetrahydroridentin B | C15 H24 O4 | -1.195 | 3.375E-03 | 267.1605 | 8.290 | 3 | DDA for preferred ion | |
| V399 | Alpha-dihydroartemisinin | C15 H24 O5 | -0.836 | 4.892E-02 | 283.1553 | 7.415 | 3 | No MS2 | |
| V363 | Cabbage identification factor 2 | C15 H12 N4 O3 S | -0.833 | 5.012E-03 | 327.0542 | 7.485 | 4 | No MS2 | |
| V362 | Carbofuran, 3OH- | C12 H15 N O4 | -0.561 | 1.735E-02 | 236.0930 | 2.850 | 3 | No MS2 | |
| V301 | Eremopetasinorol | C13 H20 O2 | -1.063 | 6.464E-03 | 207.1393 | 7.717 | 3 | DDA for preferred ion | |

| CID | Name | Formula | Fold change | p-value | m/z | RT (min) | Annotation level | MS2 | Annotation note |
|---|---|---|---|---|---|---|---|---|---|
| V283 | Gallicynoic acid F | C18 H32 O6 | -0.834 | 1.031E-02 | 343.2129 | 6.770 | 3 | No MS2 | |
| V273 | Glutamic acid glutamate | C10 H14 N2 O8 | 1.275 | 3.933E-02 | 289.0677 | 1.198 | 3 | DDA for preferred ion | |
| V165 | N-Lactoylleucine | C9 H17 N O4 | -0.659 | 9.648E-03 | 202.1088 | 1.495 | 3 | DDA for preferred ion | |
| V125 | Pantothenic acid | C9 H17 N O5 | 1.422 | 3.011E-02 | 218.1033 | 3.728 | 1 | DDA for preferred ion | confirmed by MS/MS spectral matching |
| V101 | Pluraflavin E | C36 H41 N O14 | 0.924 | 9.435E-03 | 710.2435 | 9.907 | 4 | No MS2 | |
| V96 | Prehumulinic acid | C16 H24 O4 | -0.888 | 2.763E-02 | 279.1605 | 8.330 | 3 | No MS2 | |
| V44 | Tetrahydrofurfuryl butyrate | C9 H16 O3 | -0.639 | 4.437E-02 | 171.1029 | 6.835 | 3 | DDA for preferred ion | |
| V42 | Thymidine | C10 H14 N2 O5 | 2.101 | 3.928E-02 | 241.0832 | 2.856 | 3 | DDA for preferred ion | |
| V37 | Traumatic acid | C12 H20 O4 | -0.849 | 1.327E-02 | 227.1292 | 7.485 | 3 | DDA for preferred ion | |
| V27 | Ubiquinone-2 | C19 H26 O4 | -0.991 | 7.079E-03 | 317.1761 | 8.760 | 3 | No MS2 | |
| **Extract (n = 3)** | | | | | | | | | |
| V446 | 4-O-demethylhypothemycin | C18 H20 O8 | 2.426 | 3.062E-02 | 363.1072 | 1.322 | 4 | No MS2 | |
| V278 | gamma-Glutamylisoleucine | C11 H20 N2 O5 | 0.665 | 4.520E-02 | 259.1301 | 5.419 | 3 | No MS2 | |
| V1 | γ-L-Glutamyl-(S)-2-carboxypropyl-L-cysteine | C12 H20 N2 O7 S | 2.654 | 1.526E-03 | 335.0920 | 1.412 | 3 | No MS2 | |

**Supplementary Table 5.** 10 best binding affinities, measured in binding free energy (kcal/mol) of predicted poses for all substrates, generated by AutoDock Vina in the molecular docking of SLC16A10.

| Rank | Thyroxine | Histidine | Tyrosine | Leucine | L-dopa | L-tryptophan | Phenylalanine | 3-Phenyllactic acid | Hippuric acid |
|---|---|---|---|---|---|---|---|---|---|
| 1 | -6.355 | -5.615 | -5.562 | -5.181 | -5.822 | -6.351 | -5.852 | -5.918 | -6.355 |
| 2 | -6.346 | -5.555 | -5.551 | -4.83 | -5.814 | -6.228 | -5.69 | -5.893 | -6.119 |
| 3 | -6.345 | -5.449 | -5.308 | -4.754 | -5.589 | -5.808 | -5.377 | -5.408 | -5.632 |
| 4 | -6.343 | -5.348 | -5.25 | -4.635 | -5.486 | -5.771 | -5.22 | -5.401 | -5.626 |
| 5 | -6.046 | -4.93 | -5.246 | -4.593 | -5.463 | -5.652 | -5.134 | -5.336 | -5.612 |
| 6 | -6.013 | -4.839 | -5.221 | -4.45 | -5.409 | -5.577 | -5.117 | -5.297 | -5.503 |
| 7 | -6.006 | -4.637 | -5.216 | -4.426 | -5.298 | -5.574 | -5.093 | -5.284 | -5.501 |
| 8 | -5.992 | -4.591 | -5.176 | -4.403 | -5.296 | -5.517 | -5.057 | -5.145 | -5.416 |
| 9 | -5.989 | -4.527 | -5.164 | -4.314 | -5.192 | -5.441 | -5.018 | -5.057 | -5.35 |
| 10 | -5.974 | -4.504 | -5.162 | -4.252 | -5.11 | -5.369 | -5.002 | -5.044 | -5.19 |

**Supplementary Figure 1.** Principal component analysis (PCA) of serum, serum incubated with GFP mRNA-injected *Xenopus* oocytes, and QC samples based on LC-MS/MS data data acquired in negative-ion mode (ESI-)

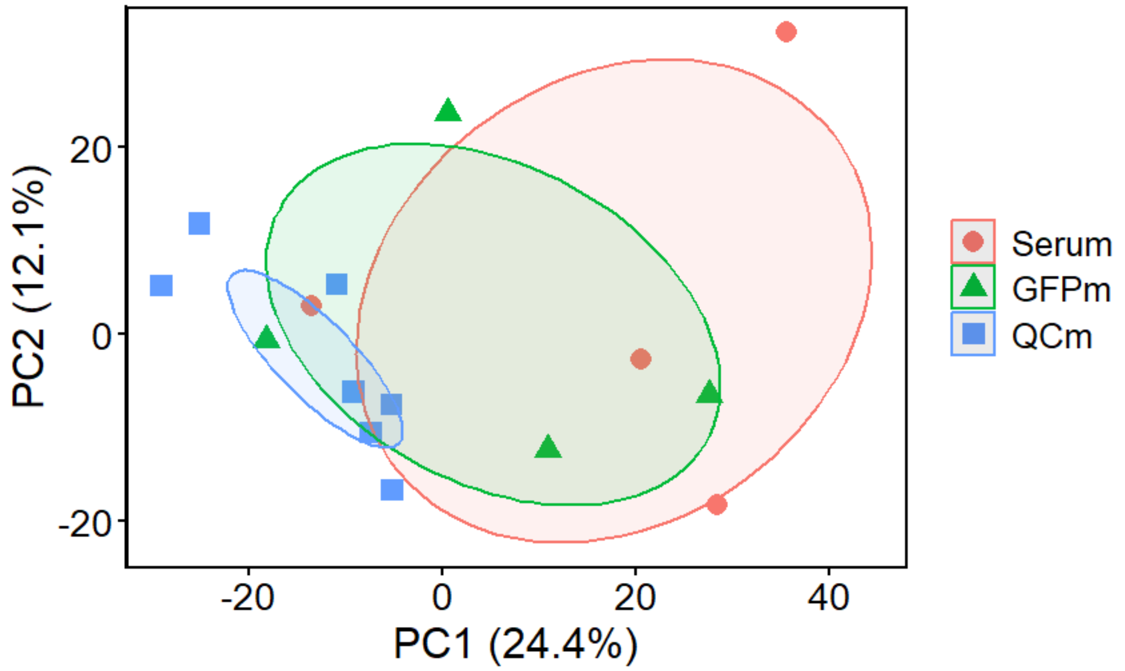

**Supplementary Figure 2.** Volcano plots showing potential metabolite exchange events based on (a) intracellular and (b) extracellular LC–MS/MS analyses data acquired in ESI- mode. Significance was defined as $p < 0.05$ and absolute $\log_2$ fold change > 0.5, indicated by the horizontal and vertical dashed lines, respectively. Data represent the average of four independent experiments, with each replicate consisting of 6–8 oocytes

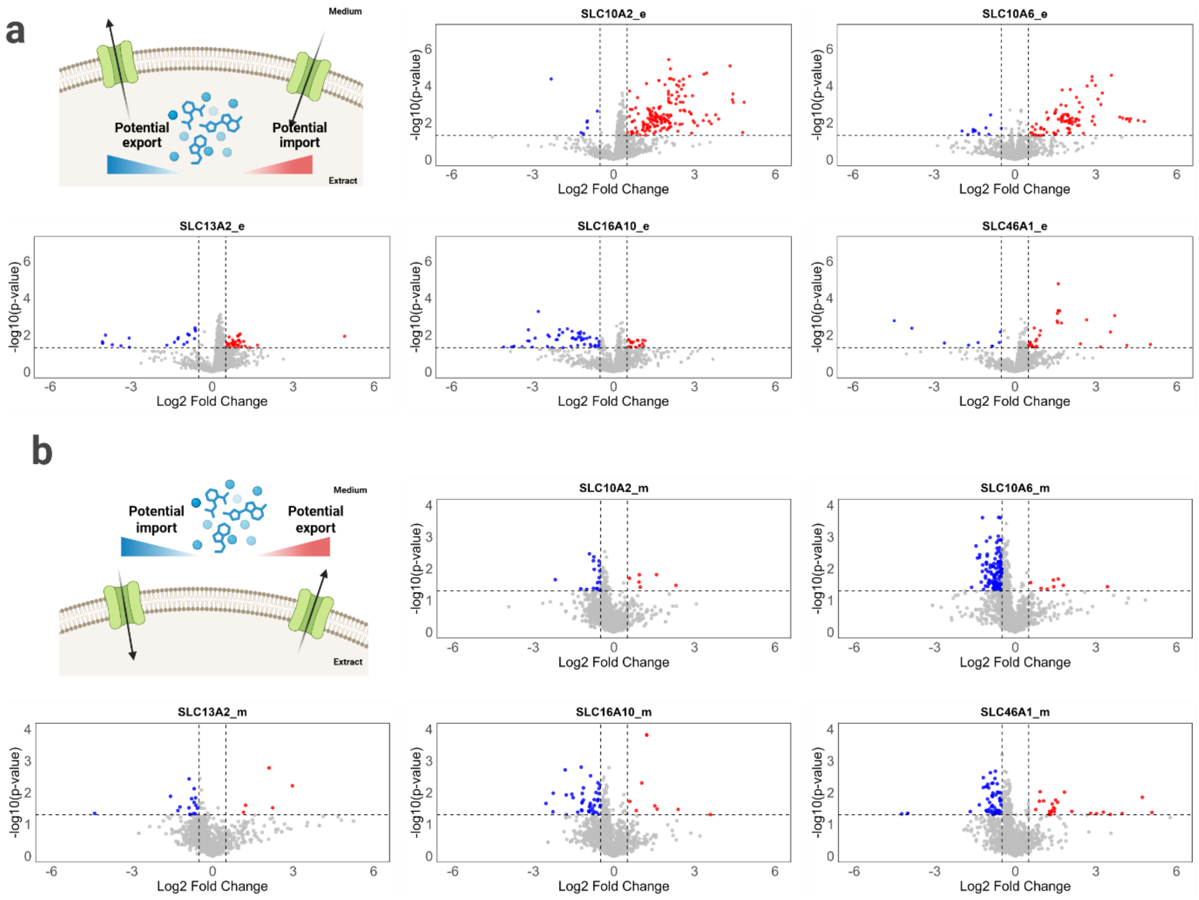

**Supplementary Figure 3.** Examples of (a) a clean LC peak (annotated as L-tryptophan) and (b) a low-quality signal (annotated as pyrimidine) that were manually inspected from the LC–MS/MS runs. Peaks were considered high quality when they showed a distinct chromatographic apex, consistent peak shape, a stable baseline with high signal-to-noise ratio, and reproducible retention time across replicates. In contrast, low-quality features lacked a clear apex and exhibited irregular or broad shapes, baseline drift, excessive noise, or unresolved spikes/shoulders, which can lead to unreliable integration and false-positive annotations

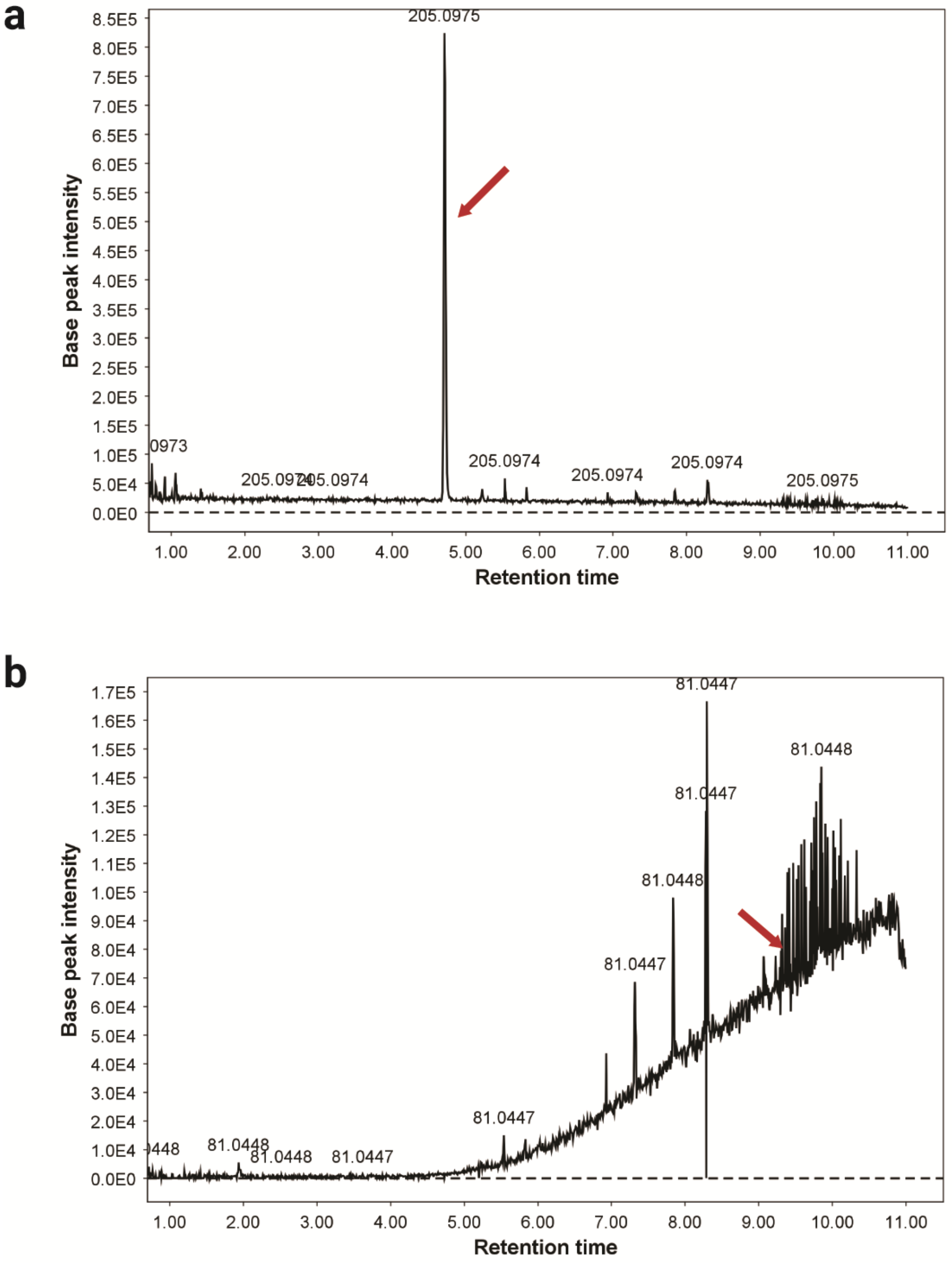

**Supplementary Figure 4.** $MS^2$ spectral alignment of glycocholic acid with an in-house reference under ESI- ionization mode. Black peaks represent the sample spectrum, and red peaks represent the in-house reference spectrum.

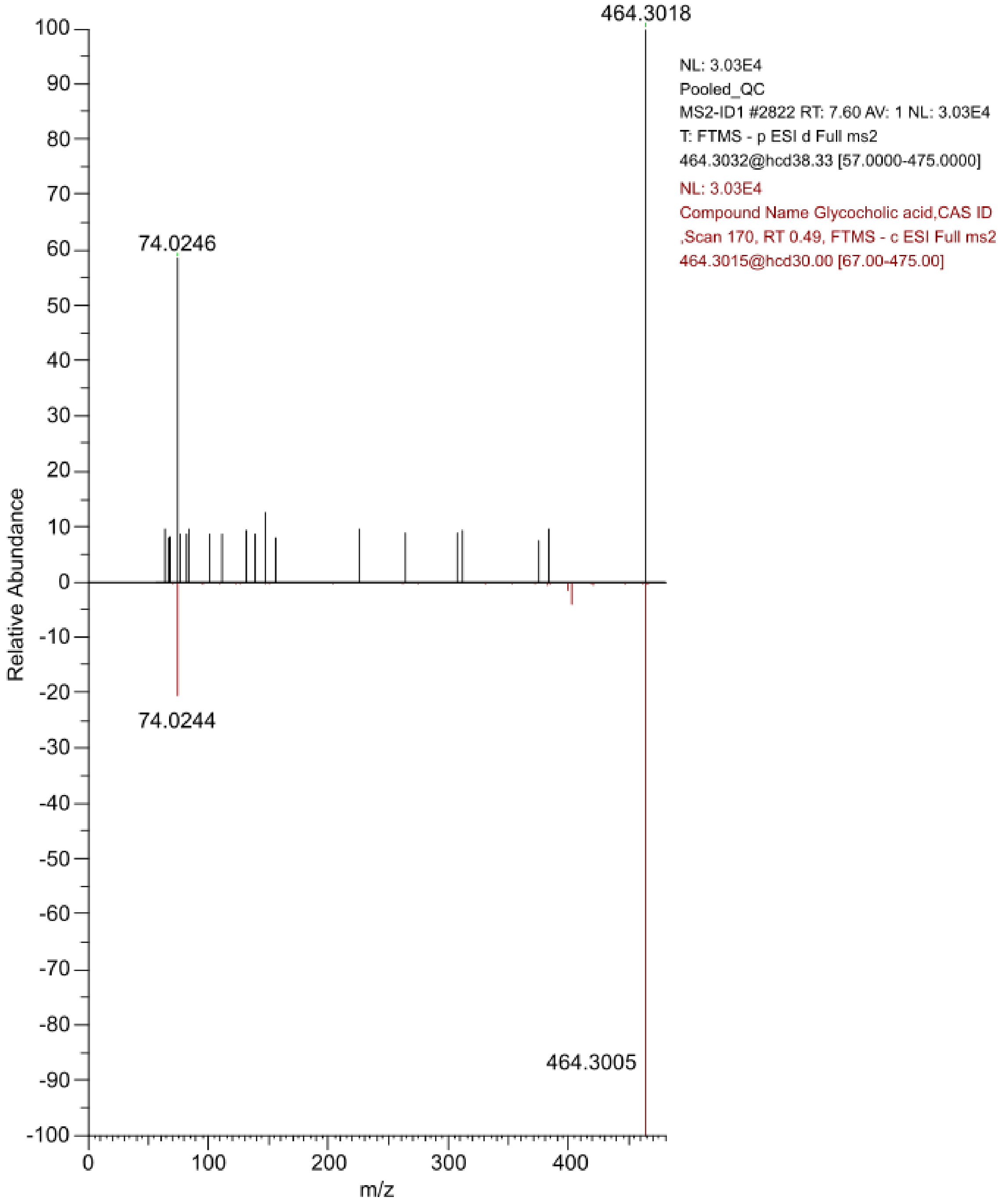

**Supplementary Figure 5.** $MS^2$ spectral alignment of L-tryptophan with an in-house reference under ESI+ ionization mode. Black peaks represent the sample spectrum, and red peaks represent the in-house reference spectrum.

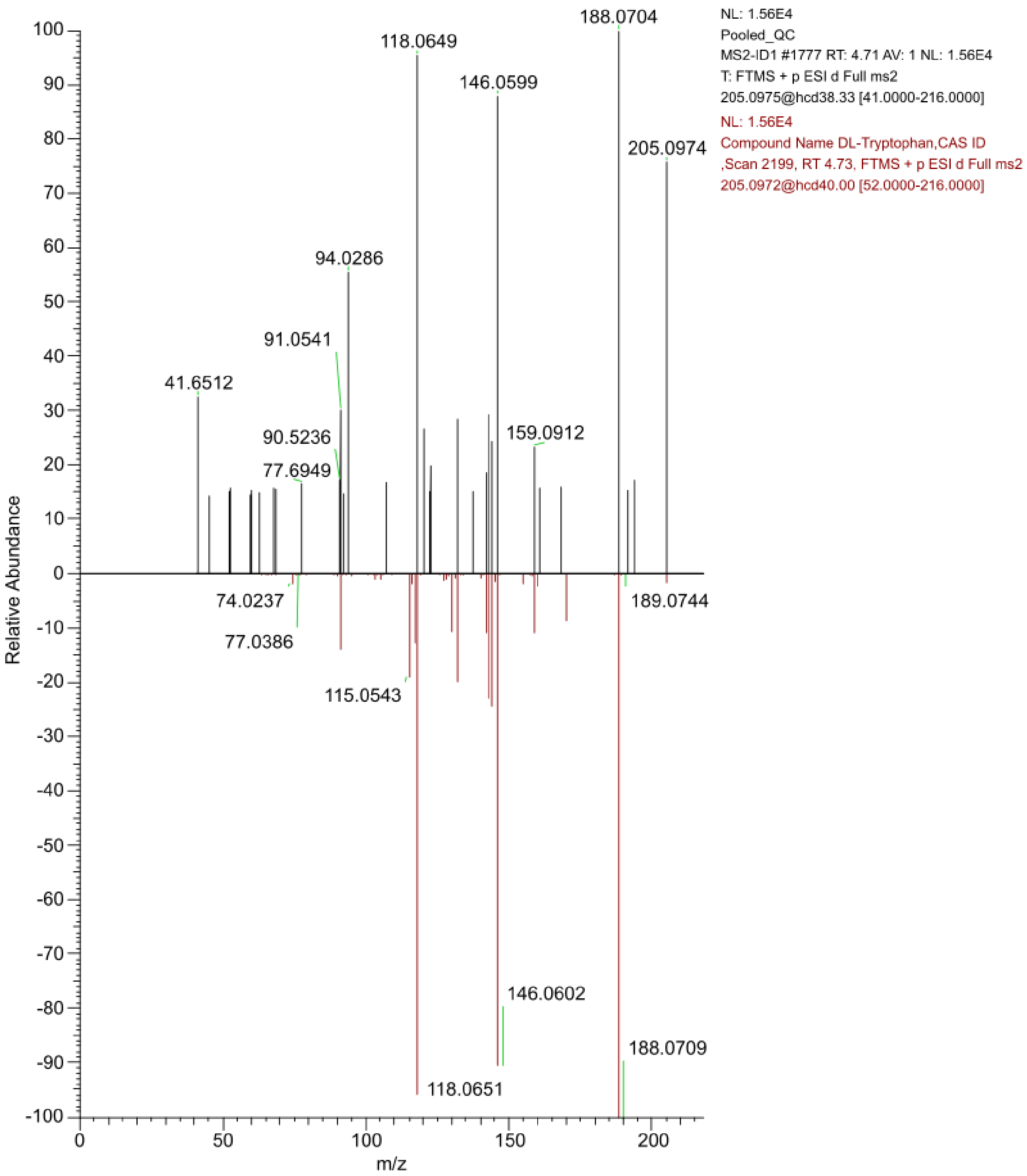

**Supplementary Figure 6.** MS² spectral alignment of L-phenylalanine with an in-house reference under ESI+ ionization mode. Black peaks represent the sample spectrum, and red peaks represent the in-house reference spectrum.

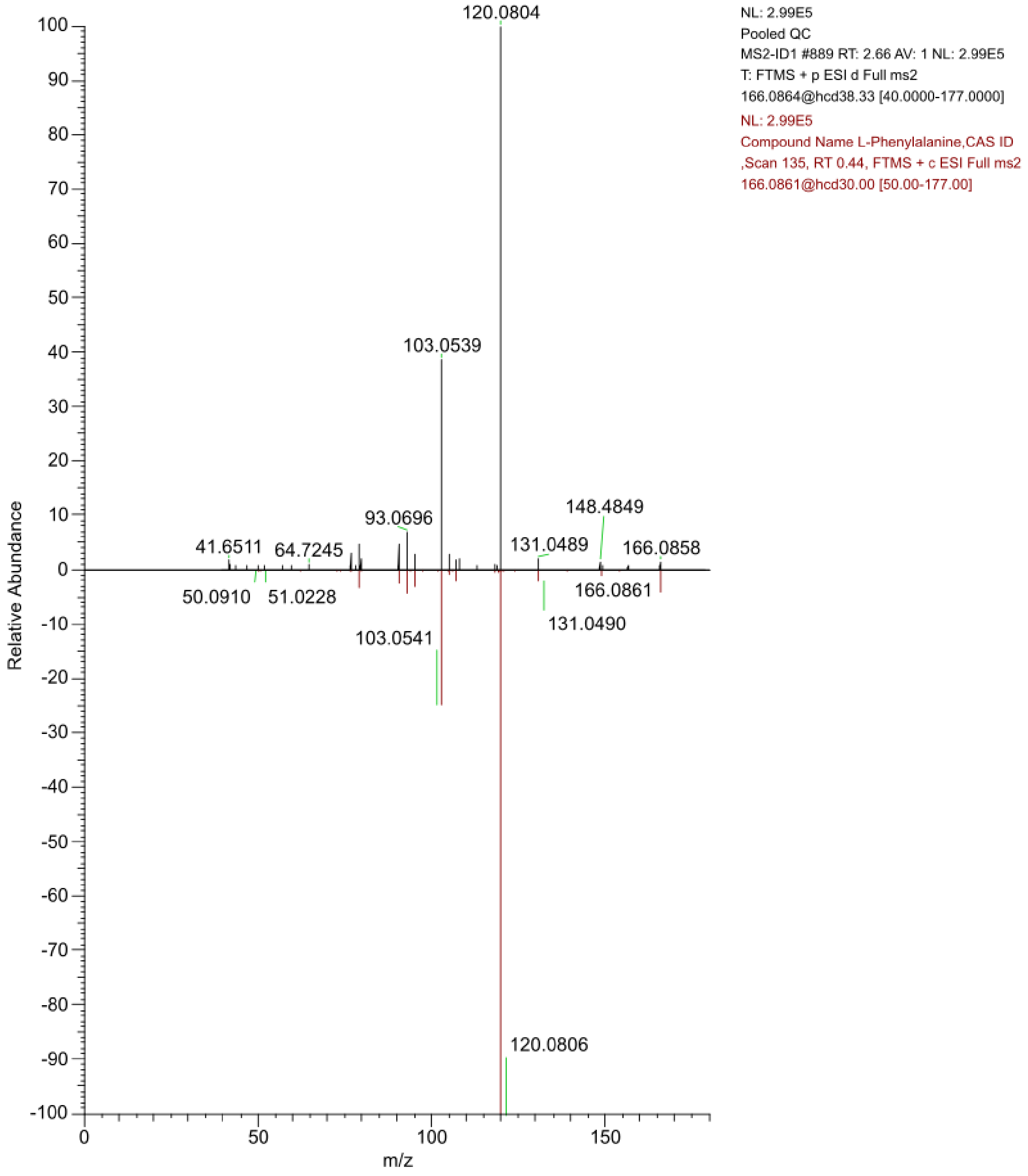

**Supplementary Figure 7.** MS² spectral alignment of L-tyrosine fragment [M+H-$NH_3$]+ with an in-house reference under ESI+ ionization mode. Black peaks represent the sample spectrum, and red peaks represent the in-house reference spectrum.

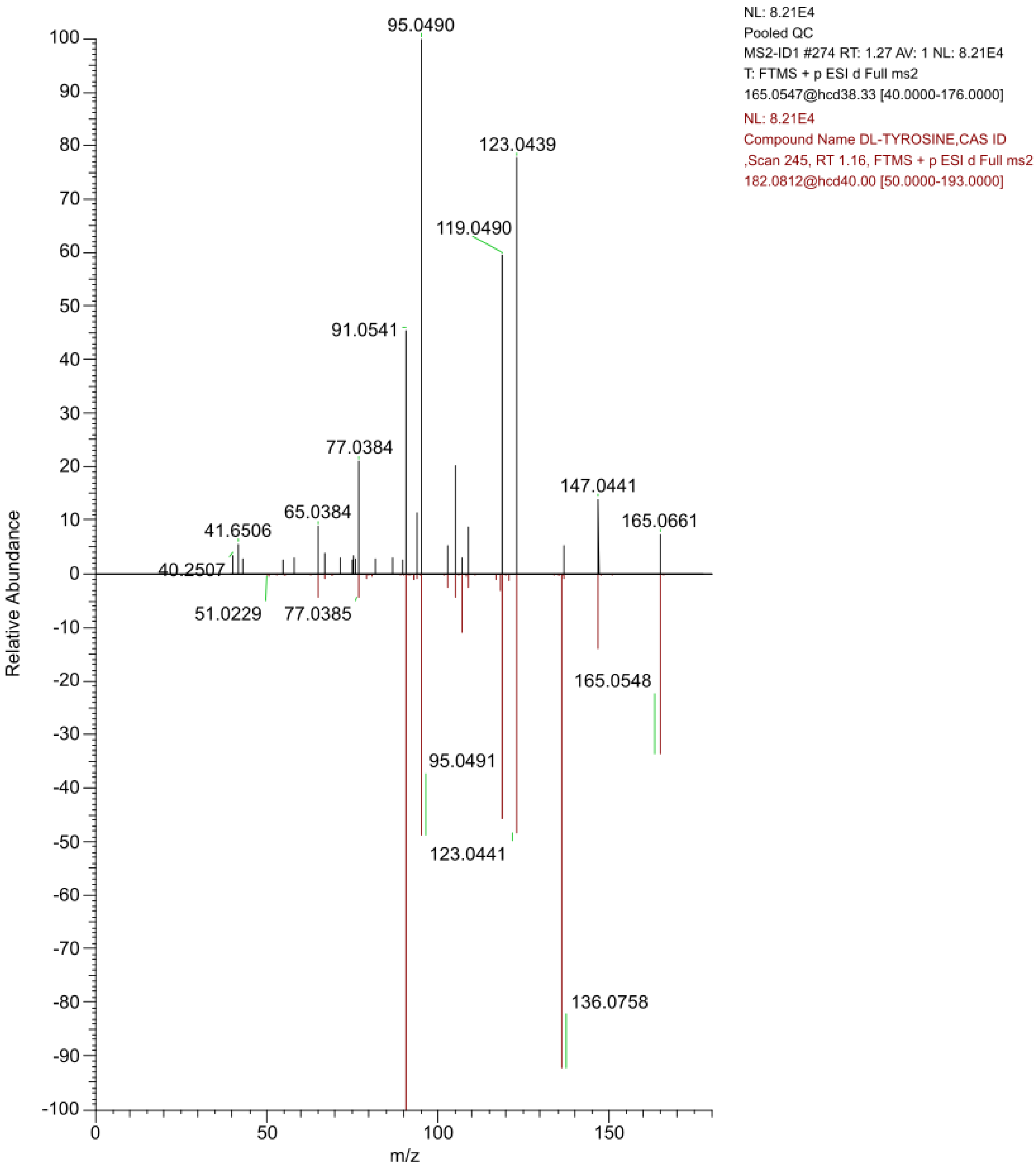

**Supplementary Figure 8.** MS² spectral alignment of hippuric acid with an in-house reference under ESI- ionization mode. Black peaks represent the sample spectrum, and red peaks represent the in-house reference spectrum.

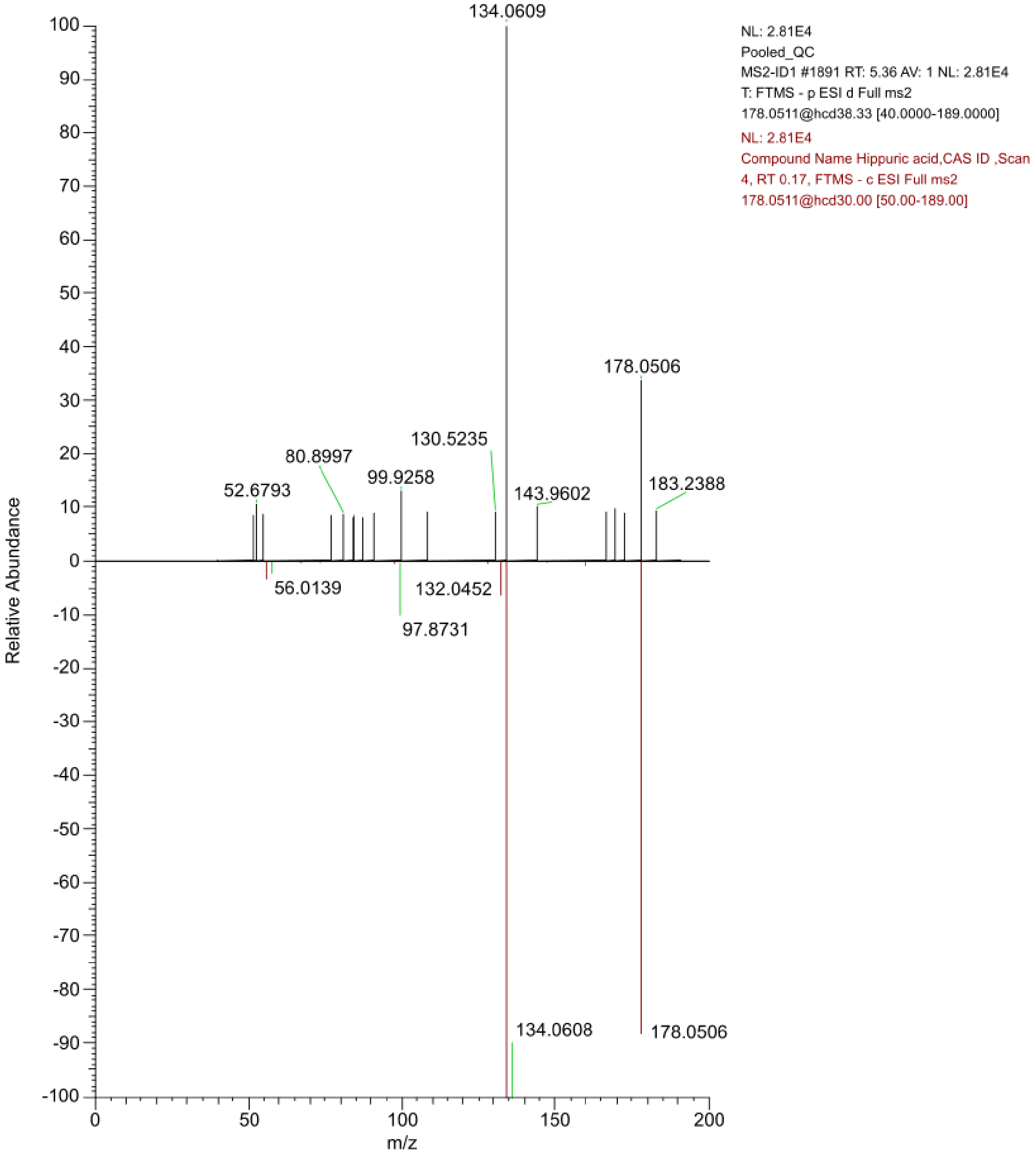

**Supplementary Figure 9.** $MS^2$ spectral alignment of 3-phenyllactic acid with an in-house reference under ESI- ionization mode. Black peaks represent the sample spectrum, and red peaks represent the in-house reference spectrum

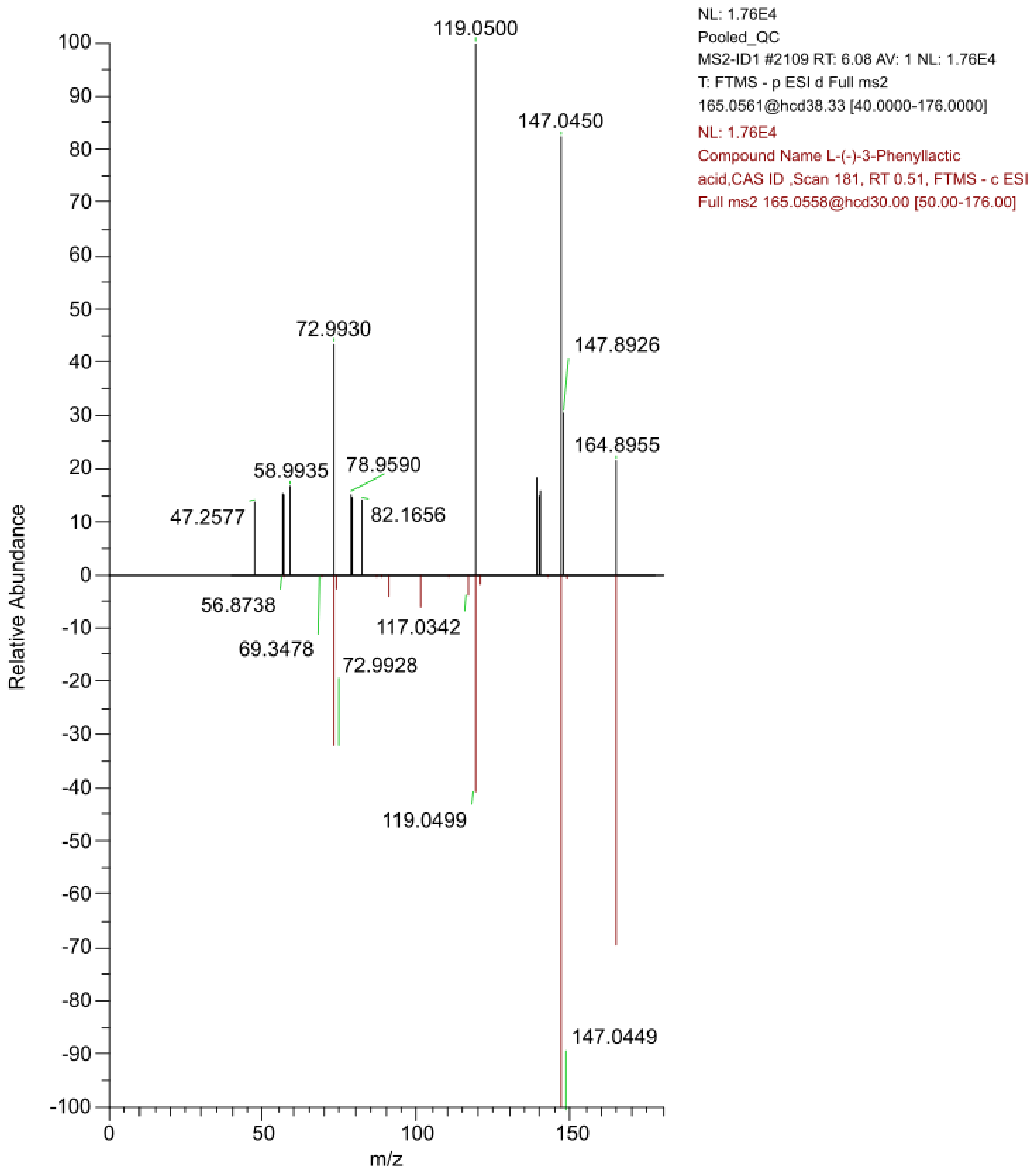

**Supplementary Figure 10.** Molecular docking results for known substrates, potential substrates and non-substrates of SLC16A10. For each substrate, the binding free energies of the top 10 poses with the best affinities, as ranked by AutoDock Vina, are shown. Data points are displayed with a dot if the centroid of the ligand pose was within the likely binding pocket, defined as the convex hull around residues that had a 3 Å distance to the ligand in the reference. Data points displayed as triangles are poses which were outside of the presumed binding pocket

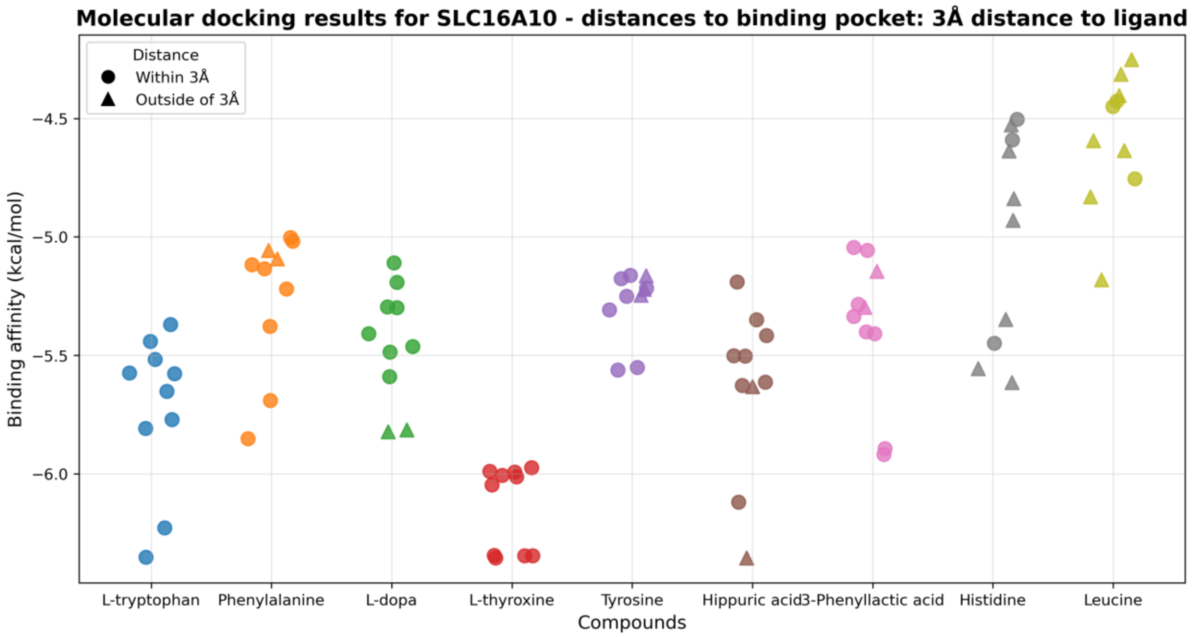

**Supplementary Figure 11.** Predicted docking poses of known and potential new substrates in SLC16A10. For each compound, the docking pose with the lowest predicted binding free energy is shown, with all structures displayed in the same orientation to allow comparison of ligand positioning within the binding region

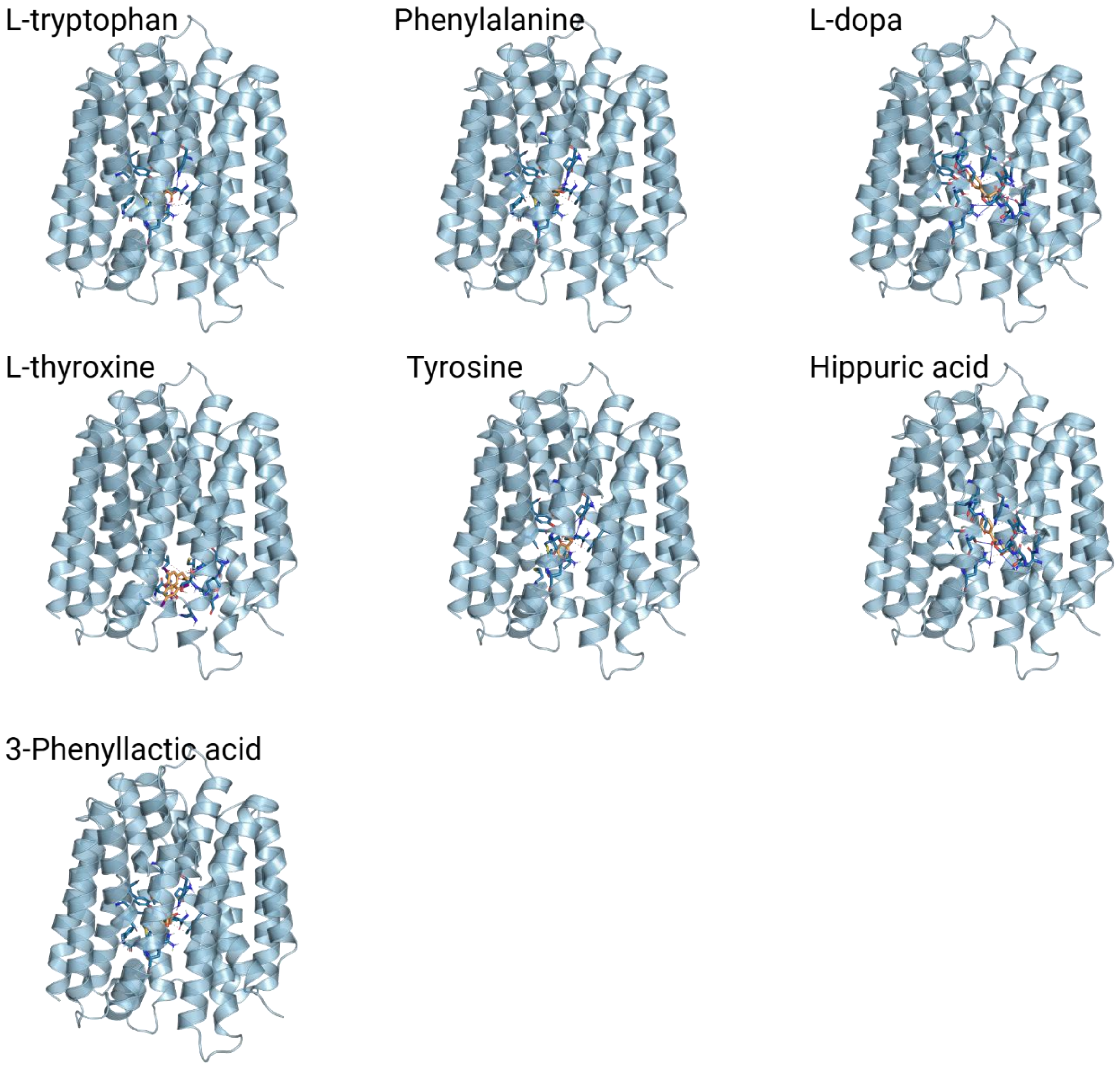
